# Fate mapping via Ms4a3 expression history traces monocyte-derived cells

**DOI:** 10.1101/652032

**Authors:** Zhaoyuan Liu, Yaqi Gu, Svetoslav Chakarov, Camille Bleriot, Xin Chen, Amanda Shin, Weijie Huang, Regine J. Dress, Charles-Antoine Dutertre, Andreas Schlitzer, Jinmiao Chen, Honglin Wang, Zhiduo Liu, Bing Su, Florent Ginhoux

## Abstract

Most tissue-resident macrophage (RTM) populations are seeded by waves of embryonic hematopoiesis and are self-maintained independently of a bone-marrow contribution during adulthood. A proportion of RTMs, however, is constantly replaced by blood monocytes and their functions compared to embryonic RTM remains unclear. The kinetics and extent of the contribution of circulating monocytes to RTM replacement during homeostasis, inflammation and disease is highly debated. Here, we identified *Ms4a3* as a specific marker expressed by granulocyte-monocyte progenitors (GMPs) and subsequently generated *Ms4a3^TdT^* reporter and *Ms4a3^Cre^-Rosa^TdT^* fate mapper models to follow monocytes and their progenies. Our *Ms4a3^Cre^-Rosa^TdT^* model traced efficiently blood monocytes (97%) and granulocytes (100%), but no lymphocytes or tissue dendritic cells. Using this model, we precisely quantified the contribution of monocytes to the RTM pool during homeostasis and inflammation. The unambiguous identification of monocyte-derived cells will permit future studies of their function under any condition.

## INTRODUCTION

Tissue-resident macrophages (RTMs) have vital roles in tissue homeostasis, inflammation and remodeling (Ginhoux and Jung, 2014), but their origins and maintenance is debated. Macrophages were originally proposed to be derived from circulating monocytes (van Furth and Cohn, 1968), but recent studies using parabiotic and fate-mapping approaches (Ginhoux et al., 2010; Guilliams et al., 2013; Hashimoto et al., 2013; Hoeffel et al., 2012; Jenkins et al., 2011; Schulz et al., 2012; Yona et al., 2013) have challenged this model, causing an important conceptual frame-shift in the field (reviewed in (Ginhoux and Guilliams, 2016)). Namely, these studies revealed that adult RTMs arise from successive waves of embryonic and adult hematopoiesis (reviewed in (Hoeffel and Ginhoux, 2015)), and the extent of contribution and the kinetics of these waves to each RTM population is tissue-specific (Ginhoux and Guilliams, 2016).

The exact contribution of adult definitive hematopoiesis to RTMs is an area of intense investigation. Using parabiosis and genetic fate-mapping approaches, Hashimoto et *al.*, showed that some RTMs (such as lung alveolar macrophages, red pulp macrophages and peritoneal macrophages) self-maintain locally throughout adult life with minimal contribution from circulating monocytes (Hashimoto et al., 2013). Others have shown that barrier tissues, such as the gut and dermis, have a notable monocyte contribution (Bain et al., 2014; Tamoutounour et al., 2013). Furthermore, bone marrow (BM)-derived monocytes can differentiate into arterial macrophages immediately after birth and locally self-renew from this point (Ensan et al., 2015), suggesting that this tissue is only temporarily “open” at birth but remains “closed” during adulthood. Adult tissues can thus be classified as: (i) closed, with no steady-state monocyte recruitment (brain, epidermis, lung and liver)/self-maintained throughout life, without or only with minimal contribution of blood monocytes; (ii) open, with fast steady-state recruitment (gut and dermis); or (iii) open, with slow steady-state recruitment (heart and pancreas) (Ginhoux and Guilliams, 2016).

The mechanisms behind these different renewal patterns are not fully understood and may be controlled by the tissue-specific microenvironment, sex and/or other factors. Bain *et al*. showed that peritoneal macrophage renewal follows a sexually dimorphic pattern, with more monocytes contributing to peritoneal macrophages in males than in females (Bain et al., 2016). The situation during inflammation is even more complicated, as partial RTM depletion occurs combined with inflammatory cell recruitment, including neutrophils and monocytes (Guilliams and Scott, 2017). These monocytes may potentially contribute to RTMs upon resolution of inflammation (Guilliams and Scott, 2017). During *Listeria monocytogenes* infection, liver-resident macrophages undergo necroptosis in conjunction with rapid monocyte infiltration to give rise to monocyte-derived Kupffer cells (KCs) (Bleriot et al., 2015). In an opposite scenario, during type 2 inflammation induced by *Litomosoides sigmodontis* infection, it is local macrophage proliferation rather inflammatory cell recruitment that controls macrophage expansion in C57BL/6 mice (Jenkins et al., 2011). These observations suggest that the developmental stage, and tissue-specific and inflammation-specific conditions, control the origins of RTMs under steady state and during inflammation. There is thus a need to precisely identify the origins of RTMs under any condition.

Monocyte fate-mapping models, including *Cx3cr1^Cre^* or *Cx3cr1^CreERT2^* (Yona et al., 2013) and *LyzM^Cre^* (Clausen et al., 1999), are not fully accurate due to the lack of lineage-specific expression of these genes used to direct Cre-recombinase expression, labeling either dendritic cells (DCs) or RTMs respectively. Here we aimed to develop a new fate-mapping mouse model specific for monocyte progenitors that could precisely measure the contribution of monocytes to RTM populations under any condition and discern monocytes from DCs. The earliest monopotent BM progenitors giving rise to monocytes are common monocyte progenitors (cMoPs) (Hettinger et al., 2013), although commitment to monocytes occurs earlier as shown by single-cell mRNA sequencing (scRNA-seq) (Giladi et al., 2018; Paul et al., 2015). cMoPs are proposed to arise from the hierarchical model of common myeloid progenitor (CMP) ➔ granulocyte-monocyte progenitor (GMP) ➔ monocyte-dendritic cell progenitors (MDP) ➔ common monocyte progenitor (cMoP) (Guilliams et al., 2018; Terry and Miller, 2014). We thus performed single-cell profiling of progenitor cells and subsequent bioinformatic analysis to identify candidate genes expressed solely by monocyte-committed progenitors to track their progeny. We identified the membrane-spanning 4-domains, subfamily A, member 3 (*Ms4a3*) gene as a specific gene to faithfully track GMPs and cMoPs but not MDPs and their DC progeny. We generated *Ms4a3^TdT^* reporter and *Ms4a3^Cre^-Rosa^TdT^* fate mapper models to specifically follow monocytes and their progenies according to different tissues, ages and sex, and to quantify their contribution to the RTM pool during homeostasis and inflammation.

## RESULTS

### *Ms4a3* is specifically expressed by monocyte-committed progenitors

We first aimed to identify a suitable gene to generate a *Cre*-recombinase-based monocyte fate-mapping model, by profiling the genes expressed in BM cMoPs and monocytes, but not in DC progenitors, such as common dendritic cell progenitors (CDPs), circulating DC precursors (pre-DCs) (to distinguish monocytes versus DCs) or differentiated macrophages (to distinguish monocyte contribution versus simple expression in macrophages). We required that this Cre driver gene to be expressed in the earliest monocyte precursors, such as GMPs or MDPs, to allow sufficient time for expression to achieve efficient recombination. MDPs are bi-potential and give rise to both DCs and monocytes (Auffray et al., 2009). Using a gene specifically expressed in MDPs could thus lead to both monocyte and DC labeling. We hypothesized, however, that within the MDP population, MDPs that are committed to the monocyte lineage co-exist with MDPs committed to the DC lineage. We thus aimed to find a gene that is uniquely expressed in MDPs that are committed to the monocyte lineage.

We performed a single-cell transcriptomic analysis of monocyte/DC progenitors by microfluidic single-cell mRNA sequencing (scRNA-seq) using the C1 Fluidigm platform **(Figure S1A for workflow**). We sorted BM cMoPs (Lin^−^CD117^+^CD115^+^CD135^−^Ly6C^+^), BM Ly6C^+^ monocytes (Lin^−^CD117^-^CD115^+^CD135^−^Ly6C^+^) and blood Ly6C^hi^ monocytes (Lin^−^CD115^+^CD11b^+^Ly6C^hi^) from wild-type (WT) C57BL/6 mice by fluorescence-activated cell sorting (FACS) (**Figure S1B and C for gating strategy**) and used scRNA-seq to generate transcriptional profiles for each individual cell (n = 38 for blood Ly6C^hi^ monocytes, n = 66 for BM cMoPs, n = 57 for BM Ly6C^+^ monocytes). We referred to our previously published dataset for MDPs (Lin^−^CD11c^−^MHCII^−^CD135^+^CD115^+^CD117^hi^), CDPs (Lin^−^CD11c^−^MHCII^−^CD135^+^CD115^+^CD117^int^) and pre-DCs (Lin^−^CD11c^+^MHCII^−^CD135^+^CD172a^-^) (Schlitzer et al., 2015). To identify putative monocyte-primed versus DC lineage-primed cells within this pool of precursors, we compared the transcriptomic signatures of each cell population to DC versus monocyte/macrophage-specific signatures (Schlitzer et al., 2015). We then performed a Connectivity Map (CMap) analysis, which is an extension of the gene set enrichment analysis (GSEA) algorithm (Lamb et al., 2006) (**Supplementary Tables 1 for gene signatures, bioinformatics analysis details see methods**). The CMap scores are scaled, dimensionless quantities that indicate the degree of enrichment or ‘‘closeness’’ of one assessed cell subset to another. Here, we identified within the whole MDP stage, single MDPs committed to the monocytic lineage versus the DC lineage (Figure 1A).

**Figure 1.**
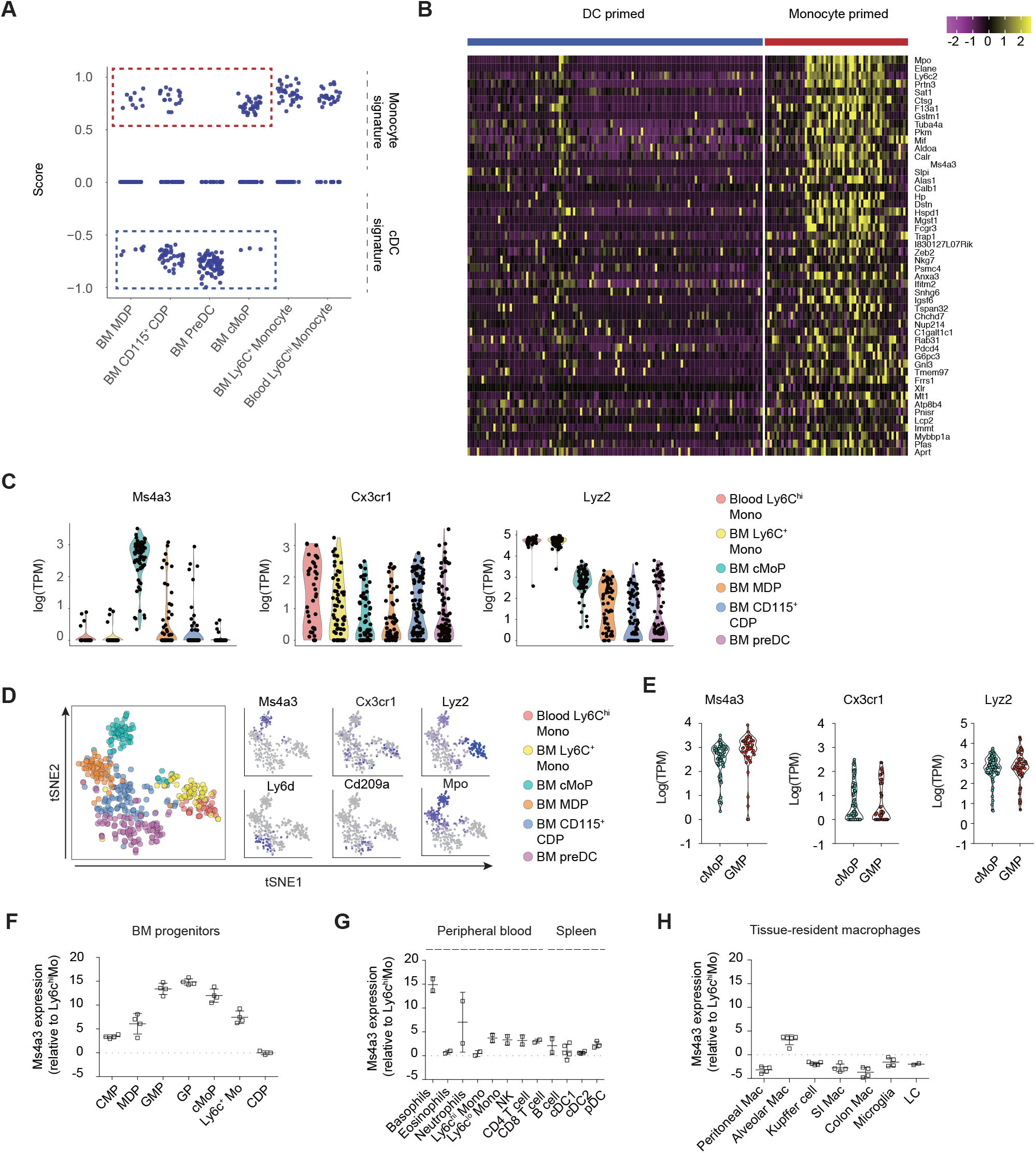
*Ms4a3* is specifically expressed by monocyte-committed progenitors. **(A)** CMap analysis of single blood Ly6C^hi^ monocytes (n = 38), BM Ly6C^+^ monocytes (n = 57), BM cMoP (n = 66), BM MDP, BM CDP and BM preDC showing enrichment for monocyte or DC signature genes. Cells with a positive CMap score were denoted as monocyte-primed cells; cells with a negative CMap score were denoted as DC-primed cells. Cells in the red rectangle (monocyte-committed cells) and blue rectangle (DC-committed cells) were used for DEG analysis. (**B)** Heatmap of the top 50 DEGs upregulated in monocyte-primed cells (indicated by the red rectangle in 1A) than in DC-primed cells (indicated by the blue rectangle in 1A), yellow indicates high expression while purple indicates low expression. (**C**) Violin plot of *Ms4a3*, *Cx3cr1* and *Lyz2* expression in blood Ly6C^hi^ monocytes, BM Ly6C^+^ monocytes, BM cMoPs, BM MDPs, BM CDPs and BM pre-DCs. Y axis is the TPM (Transcript per million reads) of Ms4a3 in each single cell. (**D**) tSNE plot of cell populations and the *Ms4a3*, *Cx3cr1* and *Lyz2* expression profiles, blue indicates high expression, grey indicates low expression. **(E)** Violin plots of *Ms4a3*, *Cx3cr1* and *Lyz2* expression in BM GMP (Lin^-^ Sca-1^-^CD117^+^CD16/32^hi^CD34^+^) and BM cMoP. (**F**) qRT-PCR analysis of *Ms4a3* expression in BM progenitors (n = 4). (**G)** qRT-PCR analysis of *Ms4a3* expression in the indicated cell types in the peripheral blood and spleen (n = 4-6). (**H**) qRT-PCR analysis of *Ms4a3* expression in RTMs in different organs. The data represent one experiment with four replicates that are representative of 2 independent experiments. The error bars represent the SEM.

As expected, progenitor populations downstream of the MDP stage mostly exhibited either monocyte/macrophage (cMoPs and monocytes) or DC (CDPs and pre-DCs) commitment. Differentially expressed genes (DEGs) between monocyte-primed progenitors versus DC-primed progenitors at the BM MDP, CD115^+^ CDP, pre-DC and cMoP stages were identified using the Seurat R package (Figure 1B). The bimodal likelihood-ratio test for single cell gene expression was used for DEG analysis, and genes with adjusted p values <0.05 were identified as being differentially expressed. We ranked the DEGs by decreasing log2 fold change, and selected the top 50 up-regulated genes in monocyte-primed progenitors compared to DC-primed progenitors (Figure 1B). After screening the expression profiles of these 50 genes in our scRNA-seq data, ImmGen (Heng et al., 2008) and bioGPS databases (Wu et al., 2009), we identified *Ms4a3* as a potential candidate gene because of its high and specific expression profile in BM monocyte progenitors (Figure 1B, C **and Figure S1D**). *Ms4a3* is a member of the membrane-spanning 4A gene family and is closely related to CD20 and the beta-subunit of the high affinity IgE receptor (FcεRIβ) (Hulett et al., 2001). Using the ImmGen database (Heng et al., 2008), we found that *Ms4a3* is highly expressed in BM GMPs, lowly expressed in MDPs (**Figure S2A**) and not expressed in DCs or various RTMs (**Figure S2B**). Exploring the bioGPS database (Wu et al., 2009), we found that *Ms4a3* to be only expressed in the BM, predominantly by GMPs (**Figure S2C**). With our single cell sequencing data, we visualized *Ms4a3* expression overlaid on dimensionality reduction with the t-Distributed Stochastic Neighbour Embedding (t-SNE) method using 7,579 genes that exhibited significantly variable expression levels between cells (Figure 1D). In this analysis, *Ms4a3* was highly expressed in cMoPs but not in CDPs, suggesting its potential utility to distinguish monocytes from DCs in a fate-mapping approach (Figure 1C and D). We also compared the expression of *Ms4a3* with *Cx3cr1* and *Lyz2*, genes used to previously make monocyte fate-mapping models. In contrast to *Ms4a3* which was only expressed in cMoPs but not in terminally differentiated blood Ly6C^hi^ monocytes, *Lyz2* was expressed at a higher level in terminally differentiated Ly6C^hi^ monocytes than in cMoPs and was also expressed by pre-DCs, while *Cx3cr1* was expressed by both monocyte-primed cells and DC-primed cells (Figure 1C and D). We also verified at the single-cell level that *Ms4a3* was expressed by GMPs (Figure 1E, **Figure S1A for workflow**). To confirm these data, we sorted BM CMPs, GMPs, granulocyte progenitors (GPs), cMoPs, MDPs, CDPs, and Ly6C^+^ monocytes (**Figure S3A for gating strategy**) and profiled *Ms4a3* expression by qRT-PCR. Consistent to the scRNA-seq data and the expression data from the ImmGen and bioGPS databases, we detected high *Ms4a3* expression in BM GMPs and cMoPs (Figure 1F). In summary, our single cell sequencing data, qRT-PCR data and expression data from public databases suggest that *Ms4a3* was highly and specifically expressed in GMPs and monocyte-committed progenitor cells and might be more suitable to specifically fate-map monocytes than *Cx3cr1* and *Lyz2*.

### Ms4a3 is not expressed by RTMs or DCs

Using Ms4a3 to build a monocyte fate mapper that could precisely measure the contribution of monocytes to RTMs and could discern monocytes from DCs requires Ms4a3 not to be expressed in mature RTMs or DCs. We first verified that Ms4a3 was not expressed by RTMs or DCs in the ImmGen and bioGPS databases (**Figure S2B and C**). To confirm these data, we sorted cell populations of interest from various tissues and profiled *Ms4a3* expression by qRT-PCR (**Figure S3B-D for gating strategy**). *Ms4a3* was expressed in basophils and neutrophils in the blood but not expressed in lymphocytes (T cells, B cells and NK cells), mature monocytes, eosinophils or splenic DCs (conventional dendritic cell 1 (cDC1), cDC2 and plasmacytoid (pDCs)) (Figure 1G). We next measured *Ms4a3* expression in several RTM populations and detected no notable expression in peritoneal macrophages, alveolar macrophages, liver Kupffer cells (KCs), gut macrophages, microglia or Langerhans cells (LCs) (Figure 1H **and Figure S3E-J for gating strategy**).

Ms4a3 is a membrane associated protein located in the peri-nuclear area but not at the cell surface (Donato et al., 2002) and no commercial antibodies for mouse Ms4a3 are available for flow cytometry limiting expression detection by flow cytometry. To confirm specific *Ms4a3* expression in GMPs and cMoPs but not DCs, RTMs or unrelated lymphoid lineages, we developed a *Ms4a3* reporter mouse by inserting an *Ires-tdTomato* sequence downstream of the *Ms4a3* stop codon (denoted as *Ms4a3^TdT^* mice hereafter) (**Figure S4**). In this model, we expected tdTomato to be expressed as a faithful reporter in cells expressing *Ms4a3*. Indeed, in the BM of *Ms4a3^TdT^* mice, the earliest tdTomato signal appeared in GMPs and was detectable in cMoPs and monocytes (albeit to a lesser extent) (Figure 2A). By contrast, high tdTomato expression was retained in BM granulocyte progenitors (Figure 2A), which is in agreement with our qRT-PCR profiling (Figure 1F). Furthermore, we confirmed high tdTomato protein expression levels in peripheral blood basophils and neutrophils, significantly less in monocytes reflecting the residual expression of the tdTomato protein initiated in the BM and no expression in lymphoid lineages (Figure 2B and C), splenic cDC1, cDC2 or pDCs (Figure 2D). These data support the concept that Ms4a3 based fate mapping models would help to distinguish monocytes from the cells of the DC lineage.

**Figure 2.**
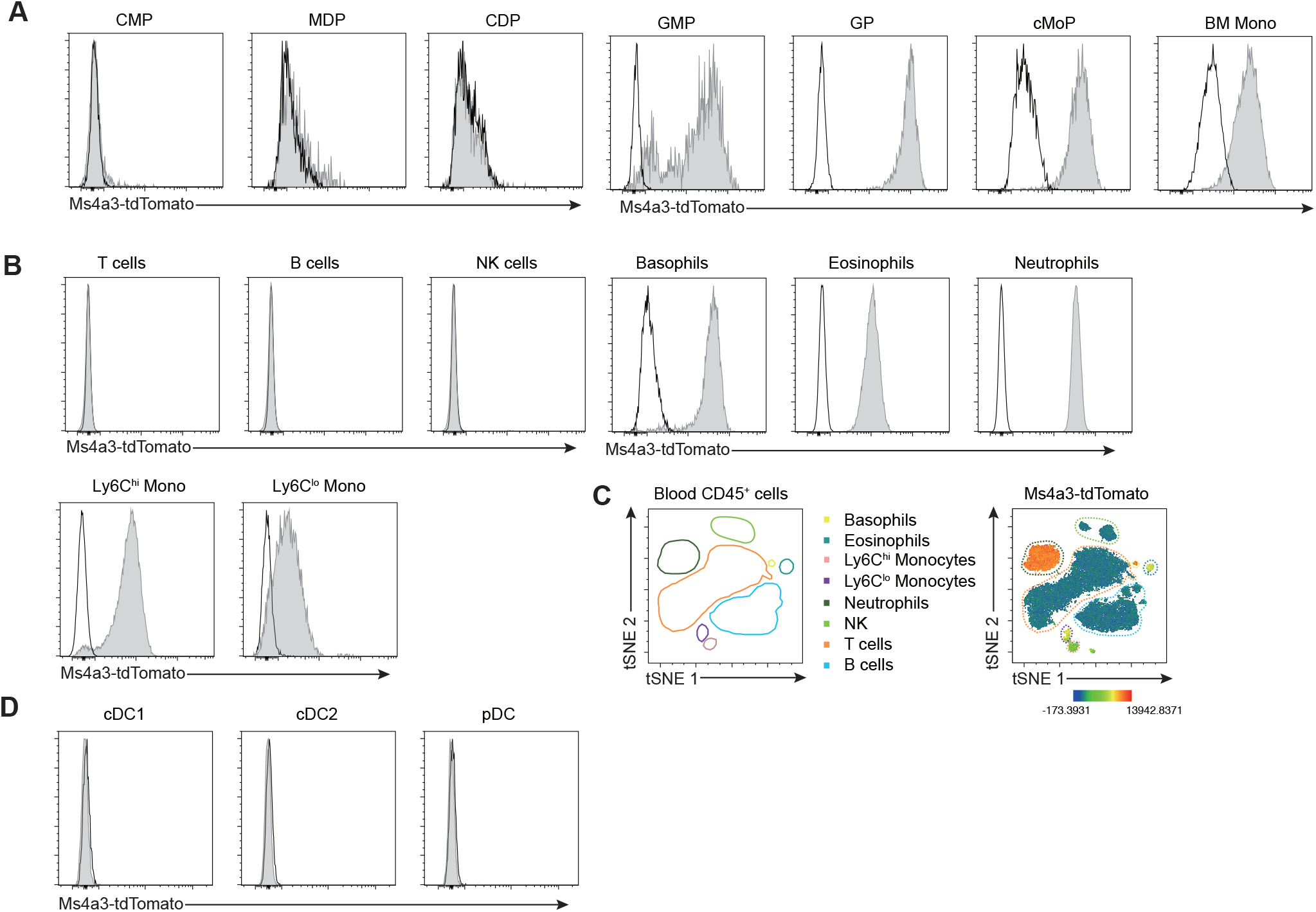
tdTomato expression in *Ms4a3^TdT^* mice. **(A)** tdTomato expression in the indicated BM cell types isolated from *Ms4a3^TdT^* (filled grey) and WT (open) mice. CMPs were defined as Lin^-^Sca-1^-^CD117^+^CD16/32^lo^CD34^+^CD135^+^CD115^-^; MDPs as Lin^-^Sca-1^-^CD117^+^CD16/32^lo^CD34^+^CD135^+^CD115^+^Ly6C^-^; CDPs as Lin^-^Sca-1^-^ CD117^lo^CD16/32^lo^CD34^+^CD135^+^CD115^+^Ly6C^-^; GMPs as Lin^-^Sca-1^-^ CD117^+^CD16/32^hi^CD34^+^CD135^-^Ly6C^-^; GPs as Lin^-^Sca-1^-^CD117^+^CD16/32^hi^CD34^+^CD135^-^ Ly6C^+^CD115^-^; cMoPs as Lin^-^Sca-1^-^CD117^+^CD16/32^hi^CD34^+^CD135^-^Ly6C^+^CD115^+^; BM monocytes as Lin^-^Sca-1^-^CD117^-^CD16/32^hi^CD34^-^CD135^-^Ly6C^+^CD115^+^. (**B**) Expression of tdTomato by indicated cell types in the peripheral blood of *Ms4a3^TdT^* (filled grey) and WT (open) mice. (**C**) tSNE plot shows the intensity of tdTomato in peripheral blood cells, the color indicates the expression intensity of tdTomato, red indicates high expression, blue indicates low expression. (**D**) Expression of tdTomato in splenic cDCs and pDCs from *Ms4a3^TdT^* mice (filled grey) and WT mice (open). Each experiment was repeated at least three times with two to three replicates and a representative plot is shown.

We next analyzed tdTomato expression in several RTM populations (brain microglia, skin LCs, liver KCs, lung alveolar macrophages, splenic macrophages, peritoneal macrophages, kidney macrophages, gut macrophages and dermal macrophages) in *Ms4a3^TdT^* reporter mice and consistent with our qRT-PCR data (Figure 1G), found that none of them expressed tdTomato (**Figure S5A-I**). Collectively, these data suggest that *Ms4a3* is a specific marker for BM GMP and cMoP stages and may be suitable for labeling GMPs and their progenies, including monocytes. Granulocytes are easily distinguished from monocytes using a granulocyte-specific marker such as Ly6G; thus, if both granulocytes and monocytes are labeled by *Ms4a3*, they can be readily identified.

### *Ms4a3^Cre^-Rosa^TdT^* model specifically and efficiently fate-maps granulocytes and monocytes

We next generated a *Ms4a3^Cre^* fate-mapper mouse model by inserting an *Ires-Cre* cassette downstream of the *Ms4a3* stop codon (**Figure S6A**) and crossed this mouse with the *Rosa26^tdTomato^* reporter strain. In the resulting *Ms4a3^Cre^-Rosa^TdT^* model, Cre recombinase is expressed in *Ms4a3*-expressing cells and recognizes its target *LoxP* sites that flank a *Stop* signal in front of *tdTomato*; the *Stop* signal is subsequently deleted, resulting in irreversible and persistent tdTomato red fluorescent protein expression in *Ms4a3*-expressing cells and their progeny (**Figure S6A**). According to the BioGPS database, where *Ms4a3* is only expressed in the BM (**Figure S2C**), we predicted no tdTomato labeling in *Ms4a3^Cre^-Rosa^TdT^* non-hematopoietic cells. Indeed, we found no tdTomato labeling in non-hematopoietic CD45^-^ cells in the brain, skin, liver, lung, kidney, pancreas, heart, salivary gland, colon or small intestine. We did, however, observe ∼60% tdTomato labeling in CD45^-^ cells in the testicles (**Figure S6B**) and confirmed low tdTomato expression in sperm by microscopy (**Figure S6C**); we did not detect any tdTomato expression in adult testicles in *Ms4a3^TdT^* reporter mice (**Figure S6D**). Going forward, to circumvent potential biases generated by germ-line recombination, we did not use *Ms4a3^Cre^-Rosa^TdT^* male mice as breeders.

We used our *Ms4a3^Cre^-Rosa^TdT^* model to first analyze tdTomato labeling in BM progenitor cells. As predicted, GMPs were the first progenitors to be highly labeled (68.7+/-1.58%) (Figure 3A, B) and almost all GMP progenies, such as cMoPs (93.5+/-0.251%) and GPs (91.8+/-0.490%) were labeled (Figure 3A, B). Reverse analysis by first gating on tdTomato^+^ cells showed that almost all tdTomato^+^ cells were CD16/32^hi^ cells, containing GMPs, cMoPs, GPs, monocytes and neutrophils (Figure 3C **and S7A**), while tdTomato^-^ cells were mainly CD16/32^lo^ cells, containing CMPs, MDPs and DC precursors (Figure 3C **and S7B**). We then analyzed tdTomato labeling in leukocyte lineages in the peripheral blood, spleen and various tissues of our *Ms4a3^Cre^-Rosa^TdT^* model. As expected, lymphoid cells in the peripheral blood, including T cells (0.03+/-0.001%), B cells (0.004+/-0.001%) and NK cells (0.120+/-0.027%), were not labeled by tdTomato, while neutrophils (99.9+/-0.048%), basophils (94.1+/-1.151%), eosinophils (99.5+/-0.309%), Ly6C^hi^ monocytes (97.3+/-0.320%) and Ly6C^lo^ monocytes (95.1+/-0.530%) were highly labeled (Figure 4A). When performing a reverse analysis by first gating on all tdTomato^+^ cells, almost all tdTomato^+^ cells were CD172a^+^CD11b^+^ myeloid cells (Figure 4B). tSNE analysis clearly showed that tdTomato^+^ cells were neutrophils, basophils, eosinophils and monocytes (Figure 4C).

**Figure 3.**
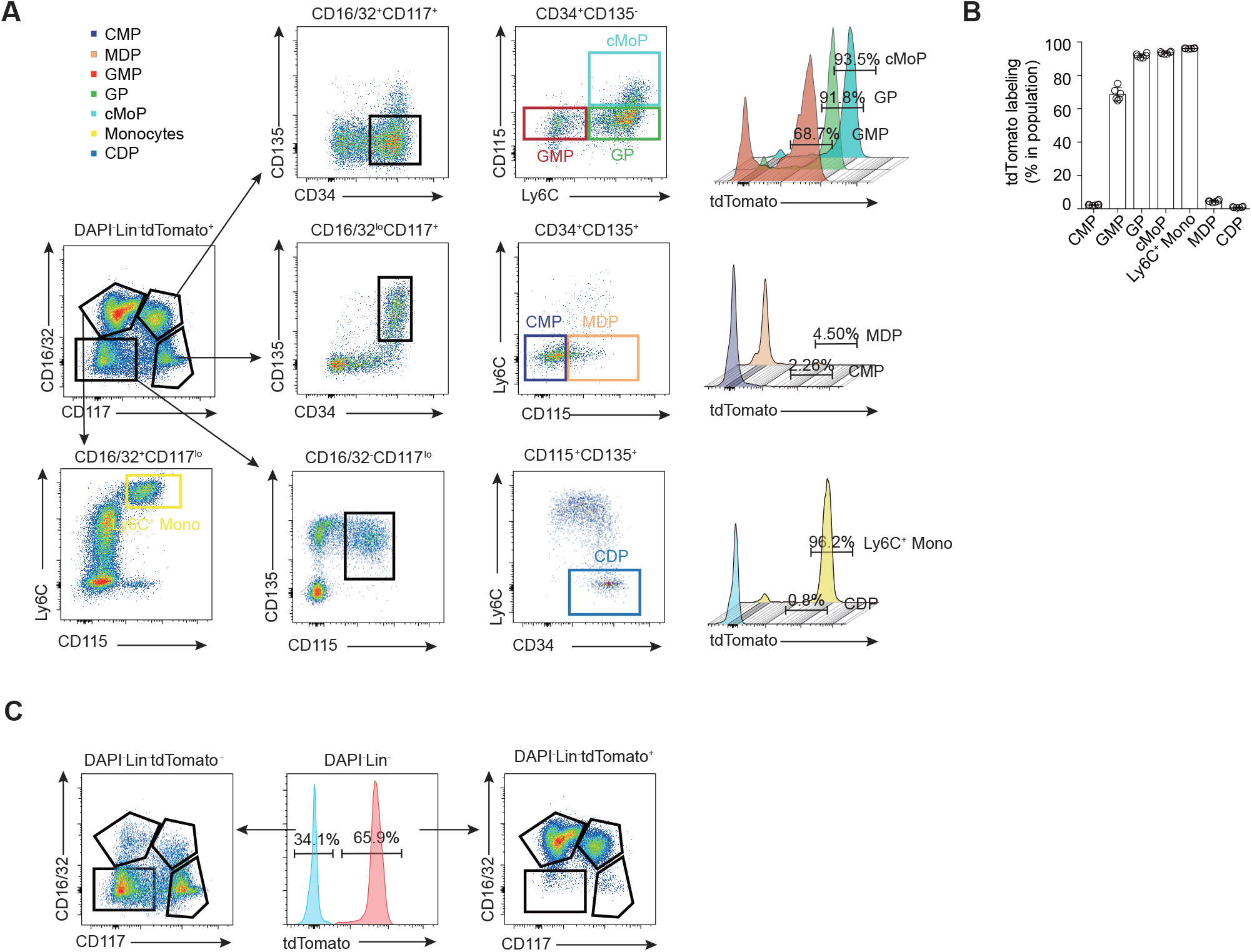
*Ms4a3^Cre^-Rosa^TdT^* labels GMPs in the BM. **(A)** Gating strategy and tdTomato labeling of BM progenitors in the BM of *Ms4a3^Cre^-Rosa^TdT^* mice. Experiments were repeated with six mice, and a representative plot is shown. **(B**) tdTomato labeling of CMP, GMP, GP, cMoP, Ly6C^hi^ monocyte, MDP and CDP in the BM of *Ms4a3^Cre^-Rosa^TdT^* mice. Data are representative of six individual mice analyzed in two independent experiments. The error bars represent the SEM. **(C)** Analysis of tdTomato^+^ and tdTomato^-^ BM cells. Plots show the distribution of tdTomato^+^ and tdTomato^-^ cells across different progenitor populations.

**Figure 4.**
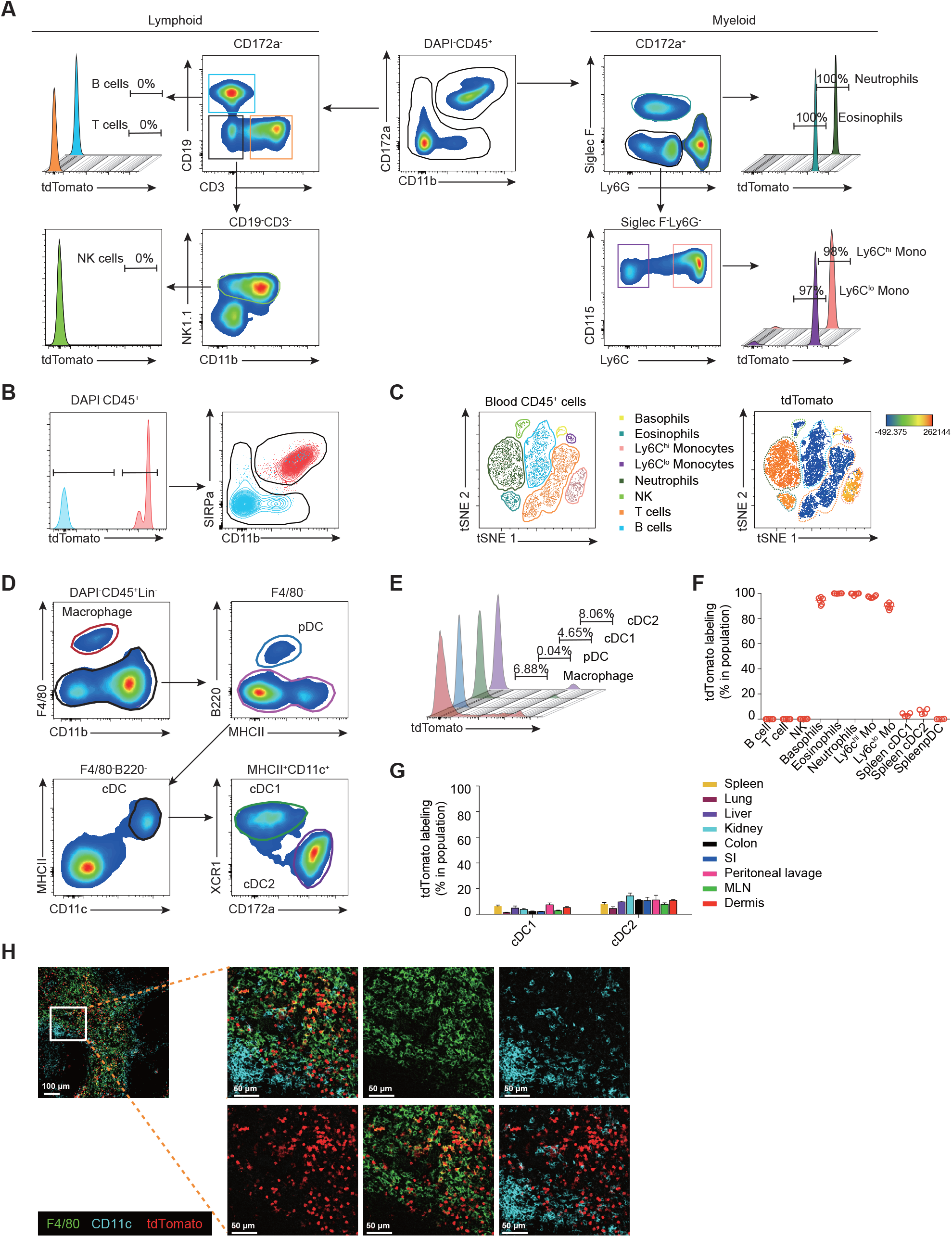
*Ms4a3^Cre^-Rosa^TdT^* model specifically and efficiently fate-maps granulocytes and monocytes. **(A**) The gating strategy and tdTomato labeling of lymphoid and myeloid lineages in the peripheral blood of *Ms4a3^Cre^-Rosa^TdT^* mice. **(B**) tdTomato^+^ cells (red) and tdtomatio^-^ cells (blue) in peripheral blood were overlaid onto CD172a vs CD11b plot. (**C**) tSNE analysis showing tdTomato expression across different cell types in the peripheral blood of *Ms4a3^Cre^-Rosa^TdT^* mice, red indicates high expression, blue indicates low expression. (**D**) Gating strategy for splenic DCs and macrophages. Experiments were repeated with six mice, and a representative plot is shown. (**E**) tdTomato labeling of splenic macrophages (CD45^+^Lin^-^F4/80^+^), pDC (Lin^-^B220^+^SIRPa^+^), cDC1 (Lin^-^ CD11c^+^MHCII^+^XCR1^+^) and cDC2 (Lin^-^CD11c^+^MHCII^+^SIRPa^+^). (**F**) tdTomato labeling in different lymphoid and myeloid lineages of *Ms4a3^Cre^-Rosa^TdT^* mice. The data are representative of 6–9 individual mice analyzed in at least two independent experiments. The error bars represent the SEM. (**G**) tdTomato labeling of cDC1 and cDC2 in different organs from 4-week-old *Ms4a3^Cre^-Rosa^TdT^* mice. Experiments were repeated twice, and the data are representative of four mice. The error bars represent the SEM. (**H**) Confocal imaging of splenic F4/80^+^ macrophages (green), CD11c^+^ DCs (Cyan), and tdTomato (red).

A *Cx3Cr1^Cre^*-based fate mapping model showed that Ly6C^lo^ monocytes are derived from Ly6C^hi^ monocytes (Yona et al., 2013); thus, labeling in Ly6C^lo^ monocytes should not be less than the labeling observed in Ly6C^hi^ monocytes. Our differential labeling pattern (95.1% versus 97.3%) suggested that Ly6C^lo^ monocytes could either be a heterogeneous population, with a minor tdTomato^-^ fraction not arising from the Ly6C^hi^ compartment, or that tdTomato^-^ cells contaminated the Ly6C^lo^ monocyte gate. To refine the gating of the Ly6C^lo^ population, we stained with the Ly6C^lo^ monocyte marker, CD43 (Ingersoll et al., 2010; Yanez et al., 2017): refining the Ly6C^lo^CD43^+^ monocyte gate resulted in tdTomato Ly6C^lo^ monocyte labeling that was identical to Ly6C^hi^ monocyte labeling (97.3+/-0.30% vs 97.7+/-0.43%, respectively) (**Figure S8A)**.

We then analyzed tdTomato labeling in splenic cDCs and pDCs. Here we detected a low level of recombination in XCR1^+^ cDC1s (3.585% +/− 0.652%) and CD172a^+^ cDC2s (6.000%+/− 1.012%), and no labeling in MHCII^int^B220^+^CD172a^+^ pDCs (0.075%+/-0.033%) (Figure 4D, E, F). We confirmed these findings by microscopy, where we observed negligible overlap between tdTomato (red) and CD11c^+^ (cyan) cells (considered DCs) in the spleen (Figure 4H). Taken together, the *Ms4a3^Cre^-Rosa^TdT^* mouse model permits faithful genetic marking of GMP progenies (monocytes and granulocytes) but not DCs.

### Monocytes do not contribute significantly to tissue DCs with age

To determine whether tdTomato labeling fluctuates with age, we analyzed peripheral blood cells and splenic DCs from *Ms4a3^Cre^-Rosa^TdT^* mice from birth up to 36 weeks. In the peripheral blood, we detected stable and efficient labeling of monocytes and neutrophils (Figure 5A **and Figure S8B**), and no labeling of lymphocytes (**Figure S8B**). In the spleen, cDC and pDC labeling did not fluctuate with age (**Figure S8C**). We used confocal microscopy to confirm tdTomato labeling of neutrophils in the spleen: all S100a9^+^ (cyan) neutrophils were tdTomato^+^ (red) (**Figure S8D**).

**Figure 5.**
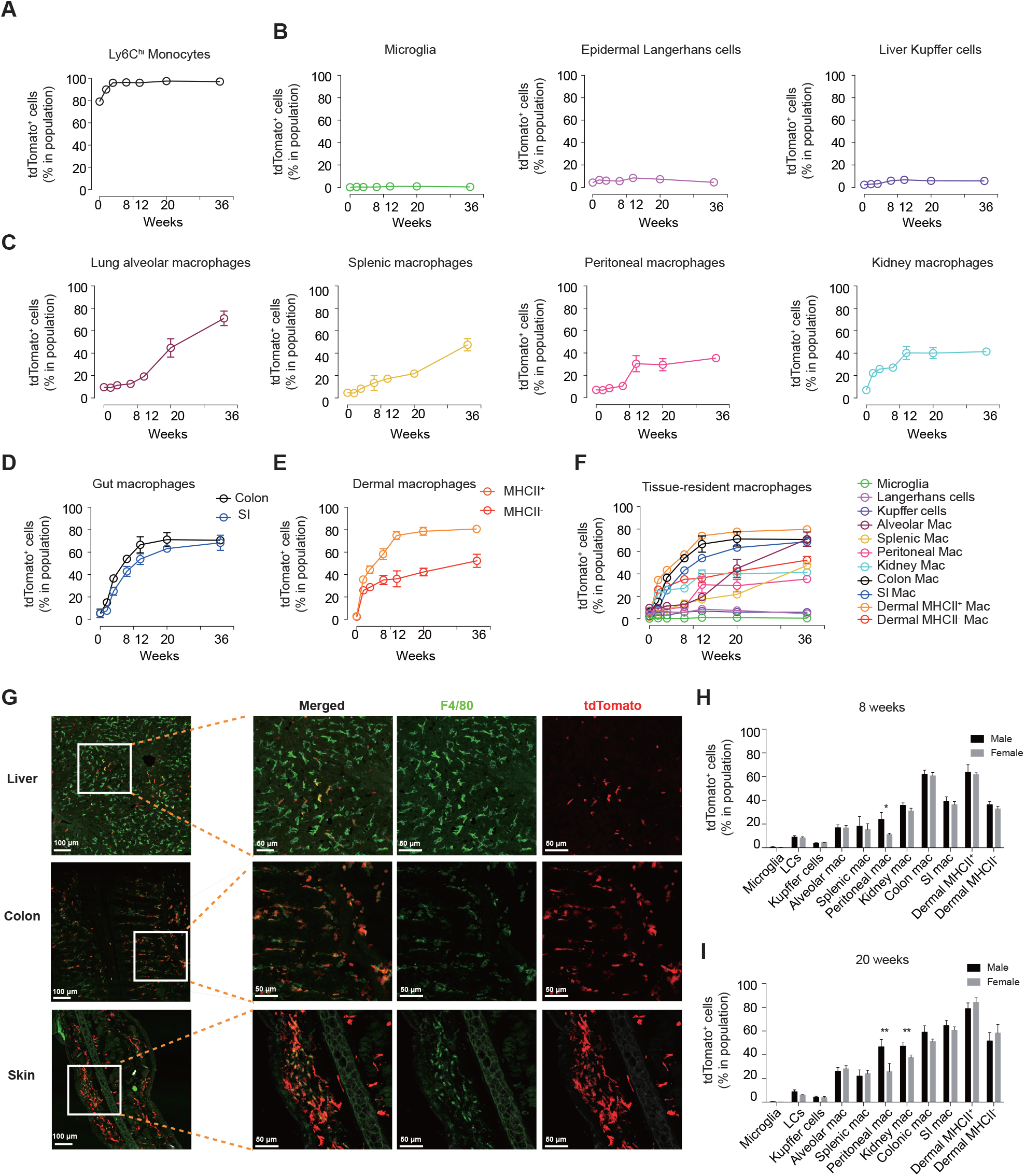
Labeling kinetics of RTMs. **(A)** Labeling kinetics of tdTomato in blood Ly6C^hi^ monocytes in *Ms4a3^Cre^-Rosa^TdT^* mice at different ages, as indicated. **(B**) Labeling kinetics of tdTomato in microglia (CD45^int^CD11b^+^F4/80^+^Ly6C^-^), epidermal Langerhans cells (CD45^+^Thy1^-^CD11b^+^F4/80^+^EpCAM^+^) and liver Kupffer cells (CD45^+^CD11b^+^F4/80^+^Tim4^+^) in *Ms4a3^Cre^-Rosa^TdT^* mice at different ages, as indicated. (**C**) Labeling kinetics of tdTomato in lung alveolar macrophages (CD45^+^SiglecF^+^CD11c^+^CD11b^lo^Ly6C^-^), splenic macrophages (CD45^+^Lin^-^F4/80^+^), peritoneal macrophages (CD45^+^CD11b^hi^F4/80^hi^) and kidney macrophages (CD45^+^CD11b^+^F4/80^+^MHCII^+^) in *Ms4a3^Cre^-Rosa^TdT^* mice at different ages, as indicated. (**D**) Labeling kinetics of tdTomato in macrophages (CD45^+^SiglecF^-^Ly6G^-^ CD11b^+^CD64^+^CD11c^-^Ly6C^-^MHCII^+^) in the colon and small intestine in *Ms4a3^Cre^-Rosa^TdT^* mice at different ages, as indicated. (**E**) Labeling kinetics of tdTomato in MHCII^+^ macrophages (CD45^+^CD11b^+^F4/80^+^CD64^+^MHCII^+^) and MHCII^-^ macrophages (CD45^+^CD11b^+^F4/80^+^CD64^+^MHCII^-^) in the dermis of *Ms4a3^Cre^-Rosa^TdT^* mice at different ages, as indicated. (**F**) Summary of the labeling kinetics of tdTomato in RTMs of *Ms4a3^Cre^-Rosa^TdT^* mice. N = 3-4 female mice analyzed per time point. (**G**) Confocal imaging showing tdTomato (red) labeling in F4/80^+^ macrophages (green) in the liver, colon, small intestine and skin and tdTomato (red). (**H**) tdTomato labeling of RTM populations from 8-week-old male (black) and female (grey) mice. n = 3 mice per sex. (**I**) tdTomato labeling of RTM populations from 20-week-old male (black) and female (grey) mice. n = 3 mice per sex. Statistical significance is indicated by, \**p*<0.05; \*\**p*<0.01. The error bars represent the SEM.

In the spleen, we found very low, but stable cDC labeling from birth up to 36 weeks, and no pDC labeling (**Figure S8C**). Using a gating strategy previously proposed to identify cDCs in murine tissues (Guilliams et al., 2016), we also showed that although tissue DCs exhibited a low level of labeling, the level was higher than in the spleen for the cDC2 in some tissues (for example dermal DC2 versus splenic DC2, 12.4 % vs 5.56 %, respectively) (Figure 4E, G **and Figure S9A**). Identifying cDC2s is notoriously less straightforward than cDC1s, as cDC2-specific markers allowing unambiguous identification have not been identified. This population may, therefore, suffer from possible contamination by monocyte-derived cells (Figure 4G). When investigating whether monocyte-derived cells could accumulate in DC populations with age (comparing 4 versus 12 weeks), we found slightly higher labeling at 12 weeks, especially in the dermis (**Figure S9B**).

These data clearly show that although the monocyte contribution to splenic and tissue DCs is minimal, the contribution of monocytes slightly increases with age in tissue cDC2. This finding highlights that care must be taken to exclude monocyte-derived cells when identifying cDC2s in non-lymphoid tissues and that our model may help to address this issue.

### The contribution of monocytes to RTMs differs under steady state between organs and shows a tissue-specific gender bias

To investigate the kinetics of RTM replenishment by monocytes in *Ms4a3^Cre^-Rosa^TdT^* mice, we analyzed tdTomato RTM labeling in various tissues (brain, epidermis, liver, lung, peritoneal cavity, kidney, spleen, colon, small intestine and dermis) at different ages (postnatal 0, 2 weeks, 4 weeks, 8 weeks, 20 weeks, 36 weeks). Of note, the labeling of blood monocytes was very efficient from birth (Figure 5A), whereas the labeling of tissue RTMs was very low in newborn mice (Figure 5B-E). This finding suggests that our fate-mapping model does not label embryonic macrophages or their precursors during development. In accordance with published reports (reviewed in (Guilliams and Ginhoux, 2016)), we found no postnatal changes in tdTomato labeling in brain microglia, epidermal LCs or liver KCs during steady-state development (Figure 5B); conversely, tdTomato labeling in macrophages slowly increased with age in the lung, spleen, peritoneal cavity and kidney (Figure 5C). As expected (Bain et al., 2016), we observed a rapid increase in tdTomato labeling in macrophages in the colon and small intestine over the first 8 weeks after birth, with a faster rate in the colon than in the small intestine (Figure 5D). Also in accordance with a previous report (Tamoutounour et al., 2013), dermal MHCII^+^ macrophages were replaced faster than MHCII^-^ macrophages (Figure 5E). We confirmed our flow cytometry-based observations (Figure 5F) by immunofluorescence analysis of the corresponding tissues (Figure 5G). Most liver F4/80^+^ cells likely to be KCs were tdTomato^-^, in agreement with their embryonic origin, while most of the gut and dermal F4/80^+^ macrophages were tdTomato^+^, underlining their derivation from tdTomato^+^ monocytes (Figure 5G).

A sexually dimorphic pattern of monocyte recruitment to the peritoneal cavity and contribution to large peritoneal macrophages (LPMs) has also been described (Bain et al., 2016), with a higher monocyte contribution to peritoneal macrophages in males than females (Bain et al., 2016). We thus compared the labeling of various RTM populations in males and females at 8 weeks and 20 weeks (Figure 5H, I). At 8 weeks, among the RTM populations analyzed, only peritoneal macrophages showed a significant difference between males and females (Figure 5H). At 20 weeks, however, we observed that peritoneal macrophages and kidney macrophages significantly differed between males and females (Figure 5I). In summary, with *Ms4a3^Cre^-Rosa^TdT^* model, we found that the contribution of monocytes to RTMs differed between organs during steady-state conditions, but also by gender in peritoneal and kidney macrophages.

### Gut macrophage subsets exhibit different monocyte contribution kinetics

Gut macrophages were previously shown to be quickly replaced by monocytes after birth (Bain et al., 2014). However, a recent study showed that gut macrophages can be divided into three transcriptionally different subsets based on Tim-4 and CD4 expression (Shaw et al., 2018). These subsets showed different turnover rates: Tim-4^+^CD4^+^ gut macrophages were locally maintained deriving from embryonic precursors, Tim-4^−^CD4^+^ macrophages had a slow turnover from blood monocytes, and Tim-4^−^CD4^−^ macrophages had a high monocyte-replenishment rate (Shaw et al., 2018). Others have suggested that Tim-4 expression marks macrophages from embryonic origin (Bain et al., 2016; De Schepper et al., 2018; Scott et al., 2016; Theurl et al., 2016). We thus decided to test the validity of these assumptions and analyze the colon and small intestine of *Ms4a3^Cre^-Rosa^TdT^* mice at early (6 weeks) and late (8 months) time points. We confirmed that monocytes contribute to Tim-4^+^CD4^+^ macrophages the least when compared to the other two subsets: at 6 weeks, tdTomato labeled 13.1+/-3.65% of Tim-4^+^CD4^+^ macrophages, 66.7+/-1.85% of Tim-4^-^CD4^+^ and 74.4+/-2.44 Tim-4^-^ CD4^-^ macrophages (Figure 6A, B). Interestingly, a clear contribution of monocytes to Tim-4^+^CD4^+^ macrophages was observed at 8 months, as indicated by an increase in tdTomato labeling to 67.0+/− 2.22%. tdTomato labeling also increased in the Tim-4^-^CD4^+^ (89.9+/-2.41) and Tim-4^-^CD4^-^ (88.9+/− 2.41%) populations. These data clearly show that Tim-4 cannot be used as a surrogate marker for embryonic origin tracing and that long-lived Tim-4^+^CD4^+^ gut macrophage populations are composed of embryonic and adult-derived cells, with proportions of the latter increasing with age.

**Figure 6.**
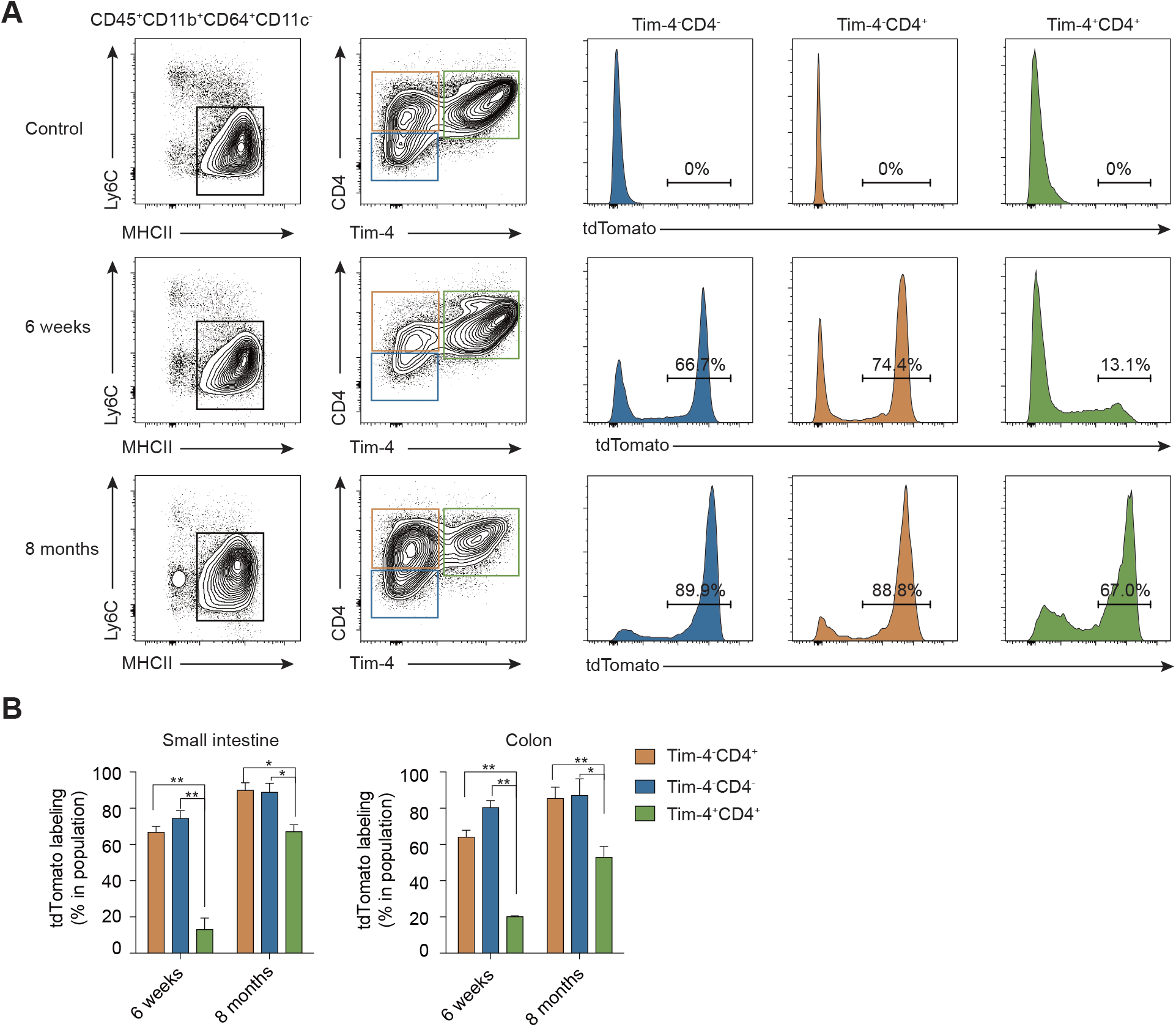
Gut macrophage subsets show different renewal kinetics. **(A)** Tim-4 and CD4 expression assessed by flow cytometry on small intestine monocytes/macrophages from WT control mice aged 6 weeks and *Ms4a3^Cre^-Rosa^TdT^* mice aged 6 weeks and 8 months old. The cells were first gated on live CD45^+^CD11b^+^CD64^+^CD11c^-^ cells, and then on macrophages (Ly6C^−^MHCII^+^). The data are representative of two independent experiments with three mice per experiment. (**B**) tdTomato labeling of three gut macrophage subsets in *Ms4a3^Cre^-Rosa^TdT^* mice aged 6 weeks and 8 months old. The data are representative of two independent experiments with three mice per experiment. Statistical significance is indicated by, \**p*<0.05; \*\**p*<0.01. The error bars represent the SEM.

### Monocyte contribution to RTMs during inflammation is model-dependent

Inflammatory stimuli often induce monocyte recruitment; these monocytes might potentially contribute to RTMs upon the resolution of inflammation. The extent of monocyte recruitment, however, is dependent on the inflammatory stimulus (Ginhoux and Jung, 2014). To validate our *Ms4a3^Cre^-Rosa^TdT^* model and investigate the contribution of circulating monocytes to RTMs during inflammation, we established various experimental scenarios of inflammation. These models included (i) thioglycollate-induced peritonitis (i.p. administration), (ii) IL-4c stimulation, (iii) LPS-induced peritonitis (i.p. administration), (iv) Lipopolysaccharide (LPS) and cytosine guanine dinucleotide (CpG) lung stimulation (i.n. administration), and (v) clodronate-loaded (CL) liposome-mediated macrophage depletion (i.v. or i.p. administration).

Thioglycollate-induced peritonitis induces peritoneal macrophage death and blood monocyte recruitment (Gautier et al., 2013). Here, we injected 1 ml of 4% thioglycollate broth into the peritoneal cavity of *Ms4a3^Cre^-Rosa^TdT^* mice and analyzed peritoneal macrophage origins after 12 h, 3 days, 3 weeks, 8 weeks and 20 weeks. At 12 h, we observed a marked decrease in the number of peritoneal macrophages, which was concomitant with an increase in the relative number of DAPI^+^ peritoneal macrophages, suggestive of induced peritoneal macrophage cell death (Figure 7A, **and Figure S10A, B**). From day 3, tdTomato labeling of peritoneal macrophages was significantly higher (74.1-92.6%) compared to the steady-state control (16.6+/-3.478%), suggesting replacement by monocytes (Figure 7B). Consistent with long-term replacement of embryonic LPMs by monocyte-derived LPMs, tdTomato labeling of peritoneal macrophages persisted at 20 weeks post-thioglycollate challenge.

**Figure 7.**
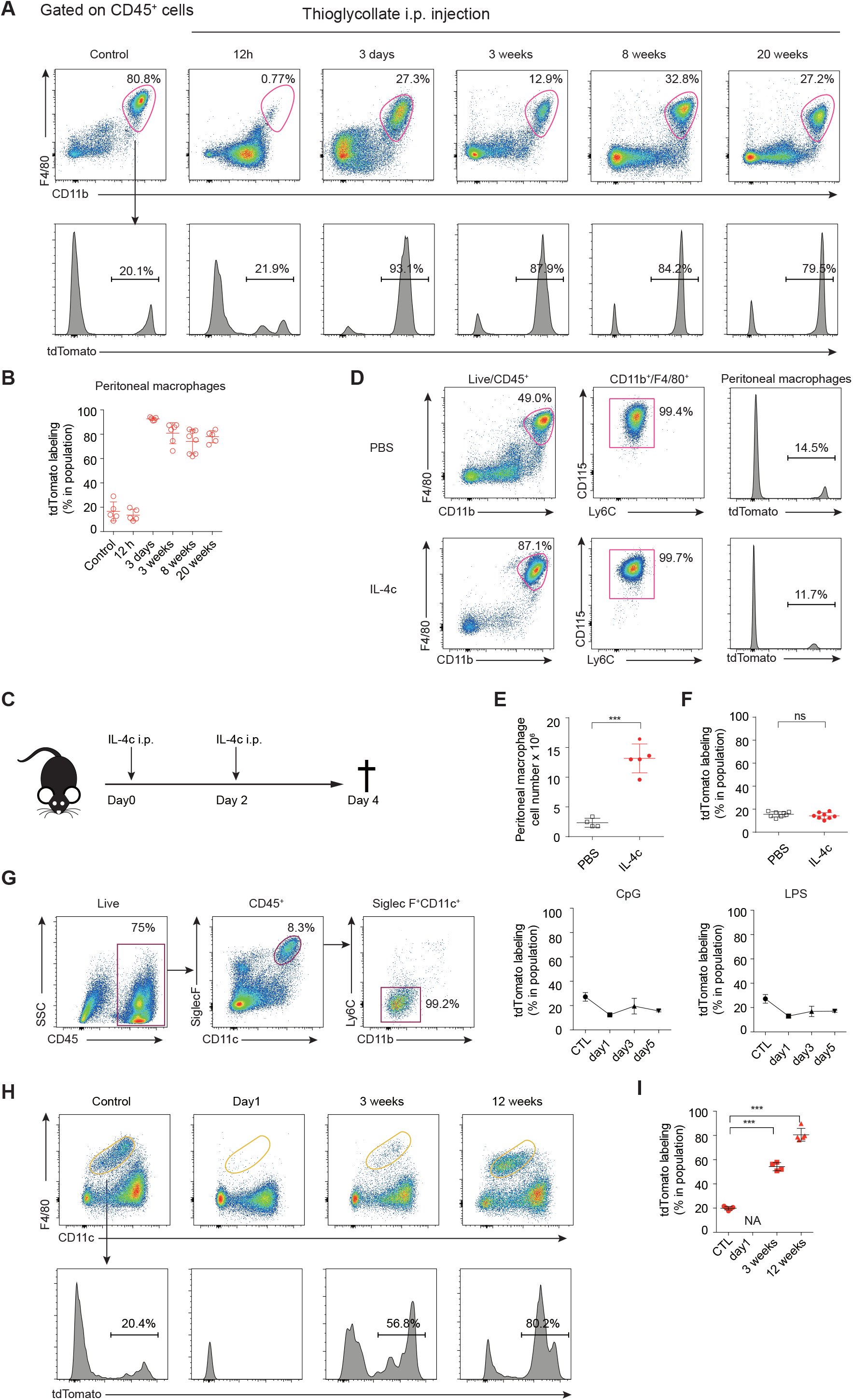
Monocyte contribution to RTMs during inflammation is model-dependent. **(A)** Flow cytometric analysis of peritoneal lavage from *Ms4a3^Cre^-Rosa^TdT^* mice at the indicated time points after i.p. injection of thioglycollate. (**B**) tdTomato labeling in peritoneal macrophages at the indicated time points after i.p. injection of thioglycollate. The data are representative of two independent experiments with 5-8 female mice per group. (**C**) Experimental protocol. Mice were injected with IL-4c on days 0 and 2, and analyzed on day 4. (**D**) Flow cytometric analysis of peritoneal lavage from *Ms4a3^Cre^-Rosa^TdT^* mice injected with IL-4c or PBS control. The data are representative of two independent experiments involving 3-4 female mice per group. (**E**) The absolute numbers of peritoneal macrophages in the peritoneal lavage isolated from female mice injected with IL-4c or PBS. The data are representative of two independent experiments involving 4-5 female mice per group. (**F**) tdTomato labeling in peritoneal macrophages from mice injected with IL-4c or PBS. The data are representative of two independent experiments involving 4-9 female mice per group. Statistical significance is indicated by \*\*\**p*<0.001; ns, not significant. The error bars represent the SEM. (**G**) Flow cytometric analysis of lung alveolar macrophages in *Ms4a3^Cre^-Rosa^TdT^* mice treated with 10 µg LPS or 50 µg CpG i.n., tdTomato labeling of alveolar macrophages was analyzed at the indicated time points, n=3-4 for each group. (**H**) Flow cytometric analysis of splenic macrophages from *Ms4a3^Cre^-Rosa^TdT^* mice at the indicated times point after i.v. injection of 200 µl clodronate liposome, n = 4-5 for each group. (**I**) tdTomato labeling in splenic macrophages from *Ms4a3^Cre^-Rosa^TdT^* mice after macrophage depletion by clodronate liposome i.v. injection, n = 4-5 for each group. Statistical significance is indicated by \*\*\**p*<0.001. The error bars represent the SEM.

We next established an opposite scenario to massive inflammatory monocyte recruitment and subsequent replacement of embryonic macrophages. Instead, we used IL-4c to induce local macrophage proliferation that is known to occur without a circulating monocyte contribution (Jenkins et al., 2011). We injected 5 µg of IL-4 combined with 25 µg of IL-4 antibody on days 0 and 2 and analyzed macrophages on day 4 (Figure 7C **and S10C**). As expected, we observed a four-fold increase in the number of peritoneal macrophages (Figure 7D, E), which was not associated with any changes in tdTomato labeling (Figure 7F). Tim-4^+^ expression did not further resolve the proliferation capacity of tdTomato^+^ versus tdTomato^-^ LPMs present before IL-4c stimulation (**Figure S10D, E**), confirming that the numbers of peritoneal macrophages increased through local proliferation of all macrophages irrespective of origins rather than by recruitment of blood monocytes.

Finally, we induced peritonitis in *Ms4a3^Cre^-Rosa^TdT^* mice by injecting LPS into the peritoneal cavity. We observed no changes in tdTomato labeling in peritoneal macrophages at any time points (days 1, 3 and 7), indicating that recruited monocytes do not differentiate into peritoneal macrophages (**Figure S10F**). It is important to note, however, that in this condition we did not observe any peritoneal macrophage cell death (**Figure S10G**). We made similar observations in other organs under inflammation. For example, we induced lung inflammation by administrating LPS or CpG intra-nasally (i.n.). In both cases, we observed no differences in tdTomato labeling of AMs between treated mice and controls on days 1, 3 and 5 post treatment, despite observing significant monocyte recruitment (**Figure S10H**). These data suggest that monocytes do not differentiate into AMs in these two inflammatory conditions (Figure 7G). Again, no significant AM cell death was observed (**Figure S10I**). Interestingly, we observed the opposite results when we fully depleted RTMs in an inflammatory context (via i.p. or i.v. clodronate liposome administration). Clodronate liposomes target and deplete various myeloid populations with phagocytic activity, including RTMs. Here, almost all macrophages in the spleen were depleted 24 h after clodronate liposomes injection (Figure 7H). By 3 weeks post-depletion, the macrophage populations that repopulated the spleen were tdTomato^+^ (control vs clodronate, 19.9+/-0.56% vs 54.3 +/− 1.70% and 80.6+/− 1.38%) (Figure 7I **and Figure S10J, K**).

Taken together, our *Ms4a3^Cre^-Rosa^TdT^* fate-mapping model shows the contribution of monocytes to RTMs under different inflammatory settings. Our data suggest that monocytes can fill the empty niche left by depleted macrophages and can develop into monocyte-derived macrophages.

## DISCUSSION

Here, we developed a neutrophil and monocyte fate-mapping model based on the high, specific expression of *Ms4a3* in BM GMPs to track the contribution of monocytes to RTMs. Using our model, we precisely quantified the contribution of monocytes to the RTM pool during steady state and inflammation. Our findings clarify the contribution of adult monocytes to RTM populations in different tissues, sexes, ages and under steady state and inflammatory conditions.

The *Ms4a* family consists of proteins with four transmembrane domains, and includes *Ms4a* members 1 (*Ms4a1*, CD20), *Ms4a*2 (FcɛRIβ) and *Ms4a*3 (HTm4 or CD20L). These proteins have key roles in regulating cell activation, proliferation and development (Eon Kuek et al., 2016). In humans, MS4A3 is 20% homologous with MS4A1/CD20 and MS4A2/FcɛRiβ (Adra et al., 1994), with the highest homology in the transmembrane domains. Importantly, mouse Ms4a3 shares 62% overall amino acid identity with human MS4A3 (Hulett et al., 2001). The first characterization of murine *Ms4a*3 showed that *Ms4a3* was only expressed in the spleen, BM and peripheral blood leukocytes at low levels (Hulett et al., 2001). A more recent report showed that *Ms4a*3 is expressed in myeloid cells (Ishibashi et al., 2018). We have extended these observations in the mouse and have precisely characterized the *Ms4a*3 expression profile first by qRT-PCR in isolated BM progenitors and mature-lineage cells from peripheral blood and splenic DCs, and then by various *in vitro* and *in vivo* assays using our *Ms4a3^TdT^* reporter mouse.

Our reporter mouse model showed that *Ms4a*3 mRNA expression is restricted to differentiating GMPs and their progeny in the BM that include neutrophils and monocytes, but also basophils and eosinophils. Interestingly, *Ms4a*3 mRNA expression is quickly lost in blood monocytes but is maintained in neutrophils. *Ms4a3* expression is also maintained in BM and blood basophils at a high level, which allows our *Ms4a3^TdT^* reporter line to be used as a specific granulocyte reporter. Similar observations for MS4A3 expression have been reported in human BM and peripheral blood cells (Ishibashi et al., 2018; Kutok et al., 2011). In the BM, MS4A3 showed a concomitant increase with progressive myeloid differentiation, whereas lymphoid or megakaryocytic-erythrocytic progenitors were entirely devoid of MS4A3 expression (Ishibashi et al., 2018). MS4A3*^+^* progenitors were shown to only produce granulocyte/macrophage colonies in *in vitro* culture (Ishibashi et al., 2018), whilst in peripheral blood, MS4A3 was highly expressed in basophils. These observations highlight the conserved MS4A3 expression profile between mouse and human and suggest its conserved functions across species.

Monocytes are proposed to arise from the hierarchical model of CMP ➔ GMP ➔ MDP ➔ cMoP ➔ monocyte (Ginhoux and Jung, 2014; Guilliams et al., 2018; Terry and Miller, 2014). However, this model has been recently challenged by Yanez *et al*., that proposed that MDPs arise directly from CMPs independently of GMPs, and that GMPs and MDPs give rise to monocytes via similar but distinct pathways through monocyte-committed progenitors (MP) and cMoPs respectively (Yanez et al., 2017). Our *Ms4a3^Cre^-Rosa^TdT^* model labelled GMPs, cMoPs and monocytes but did not label CMPs and MDPs, in agreement with the idea that MDPs do not arise from GMPs but arise from CMPs (Yanez et al., 2017). However, our MDP and cMoP labeling findings did not support the conclusion from Yanez *et al*. that MDPs gave rise to monocyte thru cMoPs while GMPs thru MPs. It is rather the contrary as cMoPs were highly labelled in the *Ms4a3^Cre^-Rosa^TdT^* model suggesting a precursor to progeny relationship, although the late acquisition of Ms4a3 expression by cMoPs cannot be formally excluded. More work need to be done to fully uncover the relationship between these differentiation stages and understand the complexity of monocyte differentiation paths.

As *Ms4a3-*specific expression starts in GMPs, it is tempting to hypothesize a unique role for this gene in cellular differentiation at this stage. Previous functional studies have shown that MS4A3 physiologically interacts with cyclin-dependent kinase (CDK)–associated phosphatase (KAP)-CDK2 complexes; this interaction enhances KAP phosphatase activity, which in turn deactivates CDK2 (Chinami et al., 2005). Fine-tuning CDK2 kinase activity may be essential for the balanced cell-cycle progression role of CDK2, which is required for the G_1_/S phase transition. Importantly, MS4A3 over-expression promotes CDK2 de-phosphorylation and leads to cell-cycle arrest at the G_0_/G_1_ phase in human myeloid U937 lymphoma cells (Chinami et al., 2005). Chinami *et al*. proposed that MS4A3 regulates the cell cycle in hematopoietic cells through its ability to control CDK2 status for the G_1_/S phase transition (Chinami et al., 2005). These results highlight an involvement of MS4A3 in controlling the cell cycle of GMP-derived cells, including monocytes and monocyte-derived cells, whereby it acts as a proliferative brake in actively dividing cells. Future studies should profile MS4A3 protein expression in blood monocytes and tissue monocyte-derived macrophages because if MS4A3 is still present in such cells once they exit the BM, the role of MS4A3 in cell-cycle regulation likely occurs in the periphery and not in the BM. We might speculate that high MS4A3 protein expression inhibits proliferation during the journey of cells migrating from the BM to the tissues. Further studies based on *Ms4a3* knockout (KO), conditional KO or over-expression in murine hematopoietic cells will help assess its functions in the BM and in tissues.

Using our *Ms4a3^Cre^-Rosa^TdT^* model, we assessed the contribution of monocytes to RTMs under steady-state conditions, and profiled several RTMs (brain, epidermis, liver, lung, peritoneal cavity, kidney, spleen, colon, small intestine and dermis) at various ages (postnatal 0, 2, 4, 8, 20 and 36 weeks). Our data are concordant with previously published reports, showing that some RTMs have no monocyte contribution (microglia, KCs and LCs) and others exhibit a tissue-specific monocyte contribution, as previously proposed (Ginhoux and Guilliams, 2016). We also observed fast (gut and dermis) versus slow (kidney, spleen and peritoneum) RTM replacement. Others have shown that even within the same tissue, macrophage subpopulations with different phenotypes exhibit different turnover rates, including in the gut (De Schepper et al., 2018; Shaw et al., 2018), heart (Dick et al., 2019) and skin (Tamoutounour et al., 2013). Here, we confirmed that in the dermis, MHCII^+^ macrophages are replaced at a faster rate than MHCII^-^ macrophages (Tamoutounour et al., 2013). These results highlight that within tissues, subpopulations of macrophages coexist with a unique homeostatic regime. The mechanisms underlying this heterogeneity could be attributed to differences in the environmental niches within the tissue or at the sub-tissular level. In agreement with this hypothesis, we recently showed that two independent monocyte-derived RTM populations coexist across tissues with distinct functional profiles in unique subtissular niches: a Lyve1^lo^MHCII^hi^CX3CR1^hi^ population that is mostly found surrounding the nerves, and a Lyve1^hi^MHCII^lo^CX3CR1^lo^ population that is often closely associated with blood vessels across tissues(Chakarov et al., 2019).

None of the RTM populations that showed a significant contribution from monocytes with time — in particular dermal MHCII^+^ and gut (of which most are MHCII^+^) macrophages — exhibited total replacement by monocytes. The level of replacement reached its asymptote, for example ∼12 weeks for dermal MHCII^+^ and gut macrophages, and no further replacement was observed thereafter. These results suggest that at its asymptotic phase, the tissue likely reaches equilibrium of monocyte recruitment, proliferation and survival/death between adult-derived RTMs and proliferation and survival/death of embryonic derived RTMs. Future studies are warranted to formally establish the exact contribution of monocyte recruitment, proliferation and survival/death at these later time points in each RTM population. Furthermore, it should be ascertained whether such equilibrium is not the result of heterogeneous populations in origin residing in different subtissular localizations, for each tissue macrophage population/subpopulation studied. Nevertheless, for more defined populations of macrophages with a known niche, our observations suggest that tissues reach equilibrium in terms of macrophage homeostasis after 12 weeks. This equilibrium might simply reflect a sign of tissue maturity in terms of niche availability — a speculation that would be in agreement with the notion that macrophage homeostasis is controlled by niche access or availability (Guilliams and Scott, 2017). The implications of these ideas are important: they suggest that if tissue maturity in terms of macrophage content is not reached before 20 weeks, the most commonly chosen 6-8 week experimental time-point used as a surrogate of adulthood may be premature.

Several studies have suggested that Tim-4 expression marks macrophages from embryonic origin (Bain et al., 2016; De Schepper et al., 2018; Scott et al., 2016; Theurl et al., 2016). Using our *Ms4a3^Cre^-Rosa^TdT^* monocyte fate-mapping model, we clearly established that while gut Tim-4^+^CD4^+^ macrophages exhibit less monocyte contribution than the other populations at 6 weeks, they do have a significant monocyte contribution at 8 months of age. These results demonstrate that Tim-4^+^CD4^+^ gut macrophage populations are composed of embryonic and adult-derived cells, with the proportion of the latter increasing with age. Consequently, we consider that Tim-4 cannot be used as a surrogate marker for embryonic origin. Although Tim-4 can be used for early time points, it is not suitable for later time points, again highlighting that murine tissues are often analyzed prematurely, when macrophage equilibrium is not reached, leading to inaccurate conclusions. Rather, Tim-4 should be considered as a marker of long-term macrophage residency in tissues (Dick et al., 2019). In a sterile model of diphtheria toxin (DT)-mediated depletion of liver-resident KCs to generate niche availability for circulating monocytes to engraft in the liver, Scott *et al*. showed that adult monocytes gradually adopt the transcriptional profile of their depleted counterparts and become long-lived self-renewing cells (Scott et al., 2016). Interestingly, while it only took several days for the engrafting monocytes to express Clec4f, a hallmark of KCs (Scott et al., 2016), monocyte-derived KCs slowly acquired Tim-4 expression, with no more than 25% of the cells expressing it 1 month after depletion. Thus, the Tim-4 expression profile does not highlight the origins of cells *per se*, but rather how much time cells have spent in their tissue of residence. The implications of such an idea are far reaching and suggest that the time spent in a tissue is crucial for cell identity, even in mature tissues considered “adult”. The mechanisms underlying such internal cellular molecular “clocks” are fascinating but remain to be established. These findings also imply that acquisition of full tissue identity takes more time than we previously thought, even in adult tissues. Furthermore, even during steady state, the monocyte-derived macrophage population will be composed of early versus late arriving cells in a continuum of differentiation. Whether certain cell functions are relevant to macrophage tissue homeostasis (such as proliferation) and can be associated with their stage of differentiation needs to be ascertained in the future. Finally, these reports all highlight the need to profile RTM populations by scRNA-seq to fully consider cellular heterogeneity.

Using our *Ms4a3^Cre^-Rosa^TdT^* mouse model, we also found the contribution of monocytes to macrophages differs according to sex and the challenge used to induce inflammation. These results highlight that the mechanisms behind different RTM renewal patterns are not fully understood and are likely controlled by many factors, including the microenvironment, age, sex and/or other factors such as the microbiome or diet; these mechanisms can now be readily tested by the scientific community using our model. For example, we observed that labeling of lung AMs increased from 10-12 weeks onwards. This finding is in contrast to previous studies that showed local replenishment of AM independent of circulating monocytes (Guilliams et al., 2013; Hoeffel et al., 2015; Schneider et al., 2014). Interestingly, the labeling of AMs differed in our Singapore facility (**data not shown**). Here, we also observed faint *Ms4a3* expression in lung AMs by PCR (Figure 1G) that could contribute to the progressive recombination in these cells, although unlikely. In addition, we found that under inflammatory conditions that deplete RTMs (such as thioglycollate-induced peritonitis), monocytes infiltrate the tissue and replenish the macrophage population, while under inflammatory conditions that do not deplete RTMs (such as LPS-induced and CpG-induced lung inflammation and LPS-induced peritonitis), we did not observe a contribution of monocytes to RTMs. These observations support that the monocyte contribution to RTMs is subject to niche access and availability (Guilliams and Scott, 2017).

In conclusion, we have generated a GMP fate mapping mouse model that permits precise and faithful identification of monocyte-derived cells during steady state conditions and disease. Using this model, we precisely quantified the contribution of monocytes to the pool of RTMs during homeostasis and inflammation and clarified the contribution of adult monocytes to various RTM populations. Our model distinguished monocytes and monocyte-derived macrophages from embryonic RTMs without cross-contamination of each population. Our model will be critical to improving our understanding of the function of embryonic RTMs and BM-derived macrophages in homeostasis and inflammation and other models of disease, from infection, cancer to metabolic diseases.

## ACKNOWLEDGEMENTS

F.G is an EMBO YIP awardee and is supported by Singapore Immunology Network (SIgN) core funding as well as a Singapore National Research Foundation Senior Investigatorship (NRFI) NRF2016NRF-NRFI001-02. This work was supported by National Natural Science Foundation of China (31470845, 81430033 and 31670896 to B.S.), Shanghai Science and Technology Commission (13JC1404700 to B.S.). The authors would like to thank Insight Editing London for editing the manuscript prior to submission.

## Author Contributions

Z.L., Y.G., S.C., C.B., X.C., A.S., W.H, R.D. and A.S. conducted the experiments; Z.L., S.C., C.B., J.C., C.A.D. B.S. and F.G. analyzed the data; Z.L. and F.G. wrote the paper; H.W. and Z.L. provided intellectual input; B.S. and F.G. supervised the project; F.G. conceptualized the study.

## METHODS

### CONTACT FOR REAGENT AND RESOURCE SHARING

Further information and requests for resources and reagents should be directed to and will be fulfilled by the Lead Contact, Florent Ginhoux.

### EXPERIMENTAL MODEL AND SUBJECT DETAILS

#### Animals

*Ms4a3^Cre^* and *Ms4a3^TdT^* mice were generated at the Shanghai Model Organisms Center, Inc. Briefly, an *Ires-Cre* or *Ires-tdTomato* gene fusion was inserted into the 3’ un-translated region (3’UTR) of the *Ms4a3* gene by CRISPR-Cas9 technique in C57BL/6 zygotes. To eliminate off-target effects, knock-in mice were then backcrossed onto a C57BL/6 background for three generations. Both strains were genotyped by PCR using the following primers:

Common forward primer 5’- AGAGAAATCATCAGGGCAGAAAT -3’;

Mutant reverse primer 5’- TTGGCGAGAGGGGAAAGAC -3’ (412 bp fragment);

Wild-type reverse primer 5’-GAAAGGGGAACAAGCGAAGAT-3’ (517 bp fragment).

*Rosa26^tdTomato^* reporter mice have been previously described (Madisen et al., 2010)All mice were bred in a specific pathogen-free animal facility at the Shanghai Jiao Tong University School of Medicine. All animal experiments were approved by the Institutional Animal Care and Use Committee (IACUC) of Shanghai Jiao Tong University School of Medicine and were performed in compliance with the University’s guidelines for the care and use of laboratory animals.

## METHOD DETAILS

### Single cell RNA-sequencing (scRNA-seq)

Cell populations (blood monocytes, BM monocytes and BM cMoPs) were isolated by flow cytometry and diluted to a final concentration range of 250–400 cells/µl. The cells were then loaded onto C1 integrated fluidic circuits (5-10-μm chip) for cell lysis, reverse transcription with oligo (dT) primers and cDNA amplification on a C1 Single-cell Auto Prep System, according to the manufacturer’s mRNA-seq protocol (Fluidigm). Array control RNA spikes were used (1, 4 and 7) (PN AM1781) according to the manufacturer’s protocol (Ambion). The cDNA generated from single cells was quantified with a QuantiT PicoGreen dsDNA Assay Kit (PN P11496; Life Technologies), and the quality was checked using High Sensitivity DNA Reagents (PN 5067-4626), according to the manufacturer’s instructions (Agilent Technologies). Only cells with high-quality cDNA were processed for subsequent library preparation. A Nextera XT Kit (PN FC-131-1096; Illumina) with dual indices (PN FC-131-1002; Illumina) was used to prepare single-cell multiplexed libraries, which were sequenced as 51-bp single-end reads on an Illumina HiSeq 2000 platform. Single-end reads were mapped to the mm9 reference genome (NCBI assembly of the mouse genome).

### scRNA-seq data analysis

CMap analysis is an extension of the GSEA algorithm (provided by the Broad Institute) in which ‘enrichment’ of a gene set (signature genes) in another gene set can be measured. CMap scores are scaled, dimensionless quantities that indicate the degree of enrichment or ‘closeness’ of one assessed cell subset to another. Monocyte and DC signature genes were identified from both the literature and our transcriptomic data, and were used as signature genes for the respective populations for CMap analysis of each single cell. The ‘enrichment’ of gene sets was tested with 1,000 permutations. Cells with a gene-expression profile that significantly correlated with signature genes were selected by a P value of <0.05 after 1,000 permutations. CMap scores were scaled to a range of –1 to 1. Cells with a positive CMap score were denoted as monocytes or monocyte primed cells, while cells with a negative CMap score were denoted as DCs or DC-primed cells.

### Quantitative Real-Time PCR (qRT-PCR)

Total RNA was isolated from sorted cells using TRIzol reagent (Invitrogen), according to the manufacturer’s protocol. Glycoblue (Invitrogen) was added as a co-precipitant when handling <10^6^ cells. cDNA was synthesized using an M-MLV First-Strand Synthesis Kit (Invitrogen, C28025-021) with oligo (dT) primers. qRT-PCR was performed using FastStart Universal SYBR Green Master with Rox (Applied Biosystems) on a ViiA 7 Real-Time PCR system (Applied Biosystems). The following primers were used for qRT-PCR:

Ms4a3 forward primer 5’- GTGGTTCTGTTTATCAGCCCTT-3’;

Ms4a3 reverse primer 5’- ACAGTGGGTAGCCTGTGTAGA-3’;

Gapdh forward primer 5’- AGGTCGGTGTGAACGGATTTG-3’;

Gapdh reverse primer 5’- TGTAGACCATGTAGTTGAGGTCA-3’.

All data were normalized to Gapdh, and quantified in parallel amplification reactions.

### Tissue preparation for flow cytometry

Blood was collected by cardiac puncture from terminally anaesthetized (isoflurane inhalation) mice; the mice were then euthanized by cervical dislocation. Peritoneal lavage was obtained by injecting 5 ml PBS containing 2 mM EDTA into the peritoneal cavity, and the washout was collected. The mice were then perfused with PBS via the left ventricle. The spleen was harvested and homogenized into a single-cell suspension using a 70 μm cell strainer and syringe plungers, then lysed in ACK lysis buffer (155 mM NH_4_Cl, 10 mM KHCO_3_, 0.1 mM EDTA) and again passed through a 70 μm cell strainer.

To obtain microglia, the brain was cut into small pieces and digested with 0.2 mg/ml collagenase type IV (C5138, Sigma) and 0.05 mg/ml DNase I (Roche) in RPM1640 medium with 10% FCS at 37°C for 60 min. The digested suspension was homogenized with a syringe with a 1.2 mm inner diameter needle. The brain-cell suspension was separated by 40%/80% layered Percoll (GE Healthcare) gradient centrifugation at 1,578 x g for 20 min at room temperature with low acceleration and no brake. The middle interface layer was collected. For newborn mice, the digested brain suspension was used for staining without Percoll separation.

For skin preparation (Tamoutounour et al., 2013), ears were split between the dorsal and ventral sections, and digested in dispase solution (Gibco) at 37°C for 90 min. The dermis and epidermis were separated and further digested as described for brain tissues. The dermis and epidermis were disrupted into single-cell suspensions using a syringe with a 1.2 mm inner diameter needle and passed through a 70 μm cell strainer.

For adult lung, liver and kidney tissues and newborn mouse tissues, tissues were cut into small pieces and digested and homogenized as described for brain preparations. Red blood cells were lysed in ACK lysis buffer.

For intestine preparation (Bain et al., 2014), the colon and small intestine were removed and washed in PBS, and the fat tissue and Peyers patches in the small intestine were removed. The intestines were opened longitudinally, cut into 0.5 cm sections and washed four times with PBS. After washing, 12.5 ml fresh calcium/magnesium-free PBS containing 5 mM EDTA and 2 mM DTT was added and the tube was incubated at 37°C with agitation for 20 min to detach the epithelial cells. The epithelial sheet was removed by vigorous shaking and the remaining tissue was washed twice with PBS, cut into small pieces and then digested and homogenized as described for brain preparations.

### Flow Cytometry

For BM progenitor analysis, BM cells were stained with APC-Cy7 conjugated anti-CD16/32 (clone 93; Biolegend) at 4°C for 15 min, and then stained with other antibodies (Supp Table 1) at 4°C for 25 min. Antibodies used for flow cytometry can be found in the **Key Resources Table**. PE-Cy7-conjugated streptavidin was used to detect biotin-labeled CD135 (clone A2F10; eBioscience). Lineage markers used in the BM progenitor analysis included CD3e, CD19, CD49b, Ly6G, Ter-119, B220, CD11c and CD11b. For flow cytometry of other samples, nonspecific antibody binding to cells was blocked by incubation with an anti-CD16/32 antibody (clone 2G8; BD Biosciences) at 4°C for 15 min, and the cells were stained with fluorophore-conjugated or biotin-conjugated antibodies at 4°C for 25 min. For intracellular staining, a commercial kit for cellular fixation and permeabilization (BD Biosciences) was used. Cells were maintained at 4 °C and analyzed on a BD Fortessa X20 or Symphony (BD Biosciences). Data were analyzed in FlowJo (FlowJo LLC). tSNE analysis was calculated with all the markers used for flow cytometry, except tdTomato.

### Cell Sorting

For peripheral blood, splenic DC and RTM cell sorting, nonspecific antibody binding to cells was blocked by incubating cells with an anti-CD16/32 antibody (clone 2.4G2; BD Biosciences) at 4°C for 15 min. The cells were then stained with fluorophore-conjugated antibodies (**Key Resources Table**) at 4°C for 25 min. FACS was performed on a BD FACS Aria III (BD Biosciences) to achieve >95% purity. Dead cells were excluded by DAPI (Invitrogen) staining.

For BM progenitor cell sorting, BM cells from the tibia and femur were used and lineage cells were depleted with a Direct Lineage Depletion Kit (Lin: CD5, CD11b, CD45R [B220], Anti-Gr-1 [Ly-6G/C], 7-4, and Ter-119; Miltenyi). Lin^-^ cells were stained with APC-Cy7 conjugated with an anti-CD16/32 antibody and incubated at 4°C for 15 min before staining with other antibodies and secondary PE-Cy7 conjugated streptavidin at 4°C for 25 min. FACS was performed on a BD FACS Aria III (BD Biosciences) to achieve > 95% purity.

### Confocal microscopy

Tissues were harvested and fixed overnight in fixation buffer containing 1% PFA. The tissues were then dehydrated in 30% sucrose before embedding in OCT freezing media (Sakura). Sections were cut to 10-µm on a Leica cryostat and blocked for 1 h at room temperature in blocking buffer containing 1% normal mouse serum, 1% BSA and 0.3% Triton X-100. The sections were stained with the indicated fluorophore-conjugated antibodies or purified antibodies overnight at 4°C in a dark, humidified chamber. Antibodies for immunofluorescence can be found in the **Key Resources Table**. A fluorophore-conjugated secondary antibody was used when necessary. Images were captured under a Leica TCS SP8 laser confocal microscope (Leica).

### Peritonitis

For thioglycollate-induced sterile peritonitis, 1 ml 4% sterile thioglycollate broth (BD Biosciences) was injected i.p. into 8-week-old female mice and the animals were analyzed at the indicated time points. For long-acting IL-4 treatment, a mix of 5 µg recombinant mouse IL-4 (Peprotech) and 25 µg anti-IL-4 mAb (clone 11B11; BioXcell, NH) was incubated for 5 min on ice to form an IL-4:anti-IL-4 complex (IL-4c). IL-4c enables sustained and slow IL-4 release. Mice were then injected i.p. with IL-4c (containing 5 µg IL-4 and 25 µg anti-IL4), or PBS vehicle control on days 0 and 2. The peritoneal lavage was analyzed at the indicated time points. For LPS-induced peritonitis, 50 µg sterile LPS (InvivoGen) was injected i.p. and peritoneal lavage was analyzed at the indicated time points.

### LPS and CpG-induced lung inflammation

Lightly anesthetized mice with isoflurane were instilled i.n. with 50 µl saline (vehicle control) or 50 µl saline containing 10 µg LPS (InvivoGen) or 50 µg CpG (InvivoGen). The mice were then analyzed at the indicated time points.

### Macrophage depletion with clodronate liposome

Mice were injected i.v. with 200 µl clodronate liposome (Yeasen) or empty liposome control to deplete RTMs in the spleen. Tissues were harvested at the indicated time points for flow cytometry and immunofluorescence staining, as described above.

## QUANTIFICATION AND STATISTICAL ANALYSIS

### Statistical analysis

The statistical analyses performed for each experiment are indicated in the figure legends. No statistical methods were used to predetermine the sample size. The experiments were not randomized. The investigators were not blinded to allocation during experiments and outcome assessment.

## SUPPLEMENTARY FIGURE LEGENDS

**Figure S1**

**(A)** scRNA-seq workflow of BM cMoPs, BM monocytes and blood monocytes using Fluidgim C1 autoprep system.

**(B)** Sorting panel for blood Ly6C^hi^ monocytes.

**(C)** Sorting panel for BM cMoPs and BM Ly6C^+^ monocytes.

**(D)** Heatmap generated with the top 10 DEGs for each population.

**Figure S2**

**(A)** *Ms4a3* expression profile in BM progenitors using ImmGen dataset.

**(B)** *Ms4a3* expression profile in DCs and RTM populations using ImmGen dataset.

**(C)** *Ms4a3* expression profile using BioGPS dataset.

**Figure S3**

**(A)** Sorting panel for BM CMPs, GMPs, GPs, cMoPs, Ly6C^+^ monocytes, MDPs and CDPs in BM.

**(B)** Sorting panel for myeloid lineage cells (basophils, neutrophils, eosinophils, and Ly6C^hi^ and Ly6C^lo^ monocytes) in peripheral blood.

**(C)** Sorting panel for lymphoid lineage cells (B cells, NK cells, CD4^+^ T cells and CD8^+^ T cells) in peripheral blood.

**(D)** Sorting panel for splenic cDCs (cDC1 and cDC2) and pDCs.

**(E)** Sorting panel for peritoneal macrophages.

**(F)** Sorting panel for lung alveolar macrophages.

**(G)** Sorting panel for liver Kupffer cells.

**(H)** Sorting panel for gut macrophages.

**(I)** Sorting panel for epidermal Langerhans cells.

**(J)** Sorting panel for brain microglia cells.

**Figure S4**

Schematic of *Ms4a3^TdT^* mice. An *IRES-tdTomato-pA* cassette was inserted after the stop codon.

**Figure S5**

**(A)** Gating strategy of brain microglia (CD45^int^CD11b^+^F4/80^+^Ly6C^-^), histogram showing the intensity of tdTomato expression in microglia of *Ms4a3^TdT^* (red) and WT (blue) mice. (**B**) Gating strategy of epidermal LCs (CD45^+^Thy1^-^CD11b^+^F4/80^+^EpCAM^+^), histogram showing the intensity of tdTomato expression in LCs of *Ms4a3^TdT^* (red) and WT (blue) mice. **(C)** Gating strategy of liver KCs (CD45^+^CD11b^+^F4/80^+^Tim4^+^), histogram showing the intensity of tdTomato expression in KCs of *Ms4a3^TdT^* (red) and WT (blue) mice. (**D**) Gating strategy of alveolar macrophages (CD45^+^SiglecF^+^CD11c^+^CD11b^lo^Ly6C^-^), histogram showing the intensity of tdTomato expression in alveolar macrophages of *Ms4a3^TdT^* (red) and WT (blue) mice. (**E**) Gating strategy of splenic macrophages (CD45^+^Lin^-^F4/80^+^), histogram showing the intensity of tdTomato expression in splenic macrophages of *Ms4a3^TdT^* (red) and WT (blue) mice. (**F**) Gating strategy of peritoneal macrophages (CD45^+^CD11b^hi^F4/80^hi^), histogram showing the intensity of tdTomato expression in peritoneal macrophages of *Ms4a3^TdT^* (red) and WT (blue) mice. (**G**) Gating strategy of kidney macrophages (CD45^+^CD11b^+^F4/80^+^MHCII^+^), histogram showing the intensity of tdTomato expression in kidney macrophages of *Ms4a3^TdT^* (red) and WT (blue) mice. (**H**) Gating strategy of gut macrophages (CD45^+^SiglecF^-^Ly6G^-^CD11c^-^CD11b^+^CD64^+^Ly6C^-^MHCII^+^), histogram showing the intensity of tdTomato expression in gut macrophages of *Ms4a3^TdT^* (red) and WT (blue) mice. (**I**) Gating strategy of MHCII^+^ macrophages (CD45^+^CD11b^+^F4/80^+^CD64^+^MHCII^+^) and MHCII^-^ macrophages (CD45^+^CD11b^+^F4/80^+^CD64^+^MHCII^-^), histogram showing the intensity of tdTomato expression in dermal macrophages of *Ms4a3^TdT^* (red) and WT (blue) mice.

**Figure S6**

**(A)** Schematic of *Ms4a3^Cre^* strategy. An *IRES-Cre* cassette was inserted after the stop codon. *Ms4a3^Cre^* mice were crossed with *Rosa^tdTomato^* reporter mice. In Cre-expressing cells, the stop codon is irreversibly removed and tdTomato expression is induced. (**B**) Flow cytometric analysis of tdTomato expression in CD45^-^ tissue cells in *Ms4a3^Cre^-Rosa^TdT^* mice. Brain, epidermis, dermis, liver, lung, kidney, pancreas, heart, salivary gland, colon, small intestine and testicular were analyzed. Experiments were repeated six times with 3-4 mice for each experiment. (**C**) Microscopic analysis showed dim tdTomato (red) expression in CD45^-^ cells in the testis of *Ms4a3^Cre^-Rosa^TdT^* mice. Cyan is CD45 and green is F4/80. (**D**) tdTomato expression in CD45-cells in testis of *Ms4a3^TdT^* reporter mice (filled grey) and WT mice (open black).

**Figure S7**

**(A)** Analysis of tdTomato^+^ BM cells. Plots show the distribution of tdTomato^+^ cells across different progenitor populations. **(B)** Analysis of tdTomato^-^ BM cells. Plots show the distribution of tdTomato^-^ cells across different progenitor populations.

**Figure S8**

**(A)** tdTomato labeling in the peripheral blood Ly6C^hi^ monocytes, Ly6C^lo^ monocytes and Ly6C^lo^ monocytes further gated on CD43^+^ (n = 9). (**B**) Kinetics of tdTomato labeling in lineages in the peripheral blood. tdTomato labeling in T cells, B cells, NK cells and neutrophils from *Ms4a3^Cre^-Rosa^TdT^* mice was analyzed at different ages. n = 3-4 mice analyzed per time point. (**C**) Kinetics of tdTomato labeling in DCs in the spleen. tdTomato labeling in cDC1, cDC2 and pDCs from *Ms4a3^Cre^-Rosa^TdT^* mice was analyzed at different ages. n = 3-4 mice analyzed per time point. (**D**) Microscopic analysis of neutrophils in the spleen showed that all S100a9^+^ neutrophils (Cyan) were positive for tdTomato (red).

**Figure S9**

**(A)** Representative flow plots of DCs in tissues using the gating strategy proposed by Guillium *et al*. (Guilliams et al., 2016). Histogram showing the tdTomato labeling in DCs in *Ms4a3^Cre^-Rosa^TdT^* mice.

**(B)** tdTomato labeling of cDC1 and cDC2 in different organs from 12-week-old *Ms4a3^Cre^-Rosa^TdT^* mice. Experiments were repeated twice, and the data are representative of four mice. The error bars represent the SEM.

**Figure S10**

**(A)** The DAPI^+^ percentage in peritoneal macrophages was analyzed at the indicated time points after i.p. thioglycollate injection. (**B**) The percentage of peritoneal macrophage in CD45^+^ cells was analyzed at the indicated time points after i.p. thioglycollate injection. Results from one experiment with three mice per time point are shown. (**C**) Images of spleens isolated from mice injected with IL-4c or PBS. (**D**) Flow cytometric analysis divided peritoneal macrophages from *Ms4a3^Cre^-Rosa^TdT^* mice into four populations with Tim-4 and tdTomato. Experiments were repeated twice and representative plots are shown. (**E**) The proportion of each macrophage populations from *Ms4a3^Cre^-Rosa^TdT^* mice. Experiments were repeated three times with n = 4 for the control and n = 3 for the IL-4c group. (**F**) tdTomato labeling of peritoneal macrophages from *Ms4a3^Cre^-Rosa^TdT^* mice injected with LPS was analyzed at the indicated time points, n=3-4 for each group. (**G**) DAPI^+^ percentage in peritoneal macrophages was measured at the indicated time points after i.p. administration of LPS. Results from one experiment with three mice per time point are shown. (**H**) The percentage of monocytes (blue) and neutrophils (orange) in live cells at the indicated time points after i.n. administration of CpG or LPS. Results from one experiment with three mice per time point are shown. (**I**) DAPI^+^ relative numbers in alveolar macrophages was analyzed at the indicated time points after i.n. administration of CpG or LPS. Results from one experiment with three mice per time point are shown. (**J**) Flow cytometric analysis of splenic macrophages from *Ms4a3^Cre^-Rosa^TdT^* mice at the indicated time points after i.p. injection of 200 µl clodronate liposome and (**K**) tdTomato labeling in splenic macrophages from *Ms4a3^Cre^-Rosa^TdT^* after macrophage depletion by clodronate liposome i.p. injection, n= 5-6 for each group. Statistical significance is indicated by \*\*\**p*<0.001. The error bars represent the SEM.

