## Supplementary figures and images for "Fate mapping via Ms4a3 expression history traces monocyte-derived cells"

### Supplementary Figures

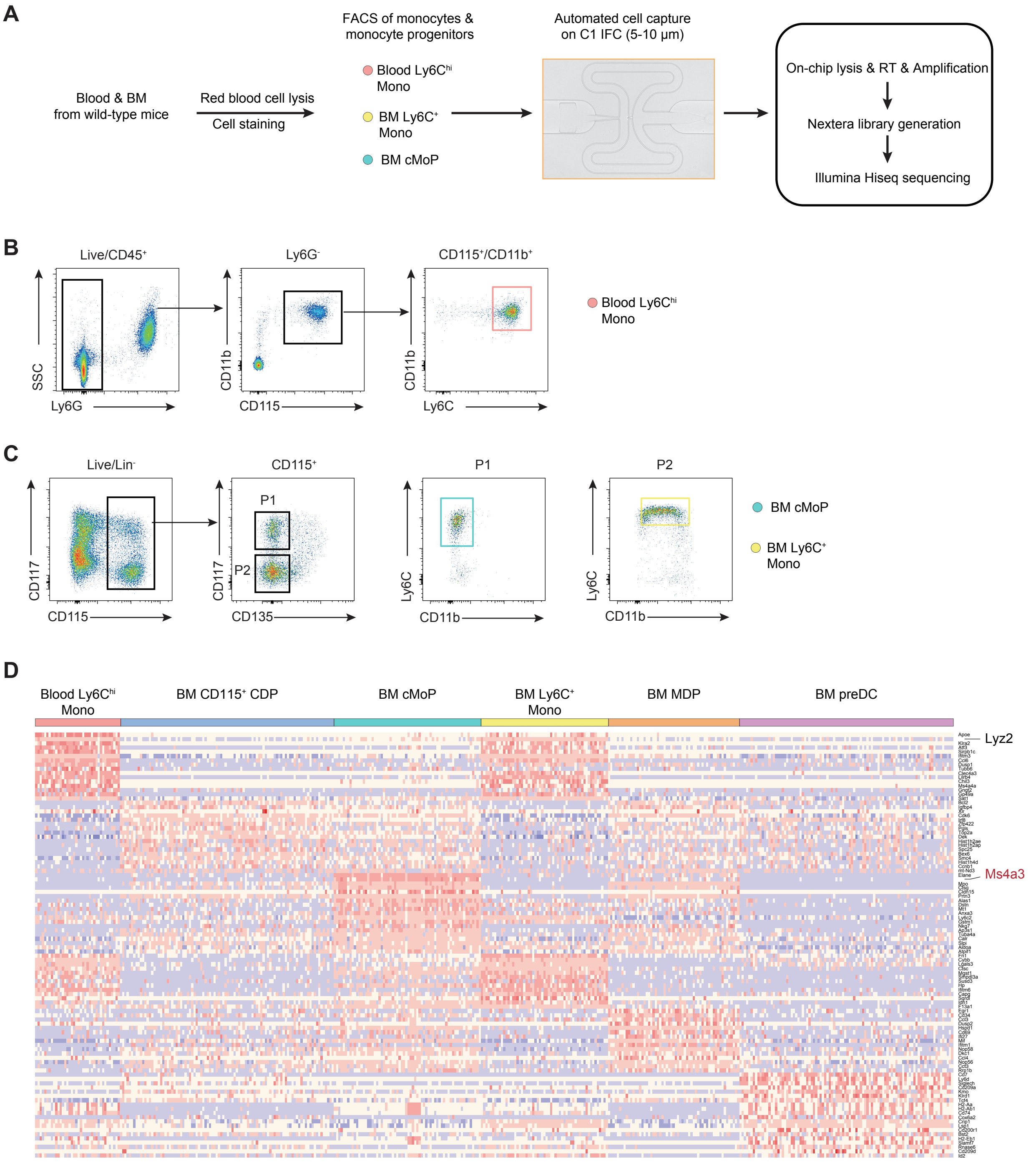

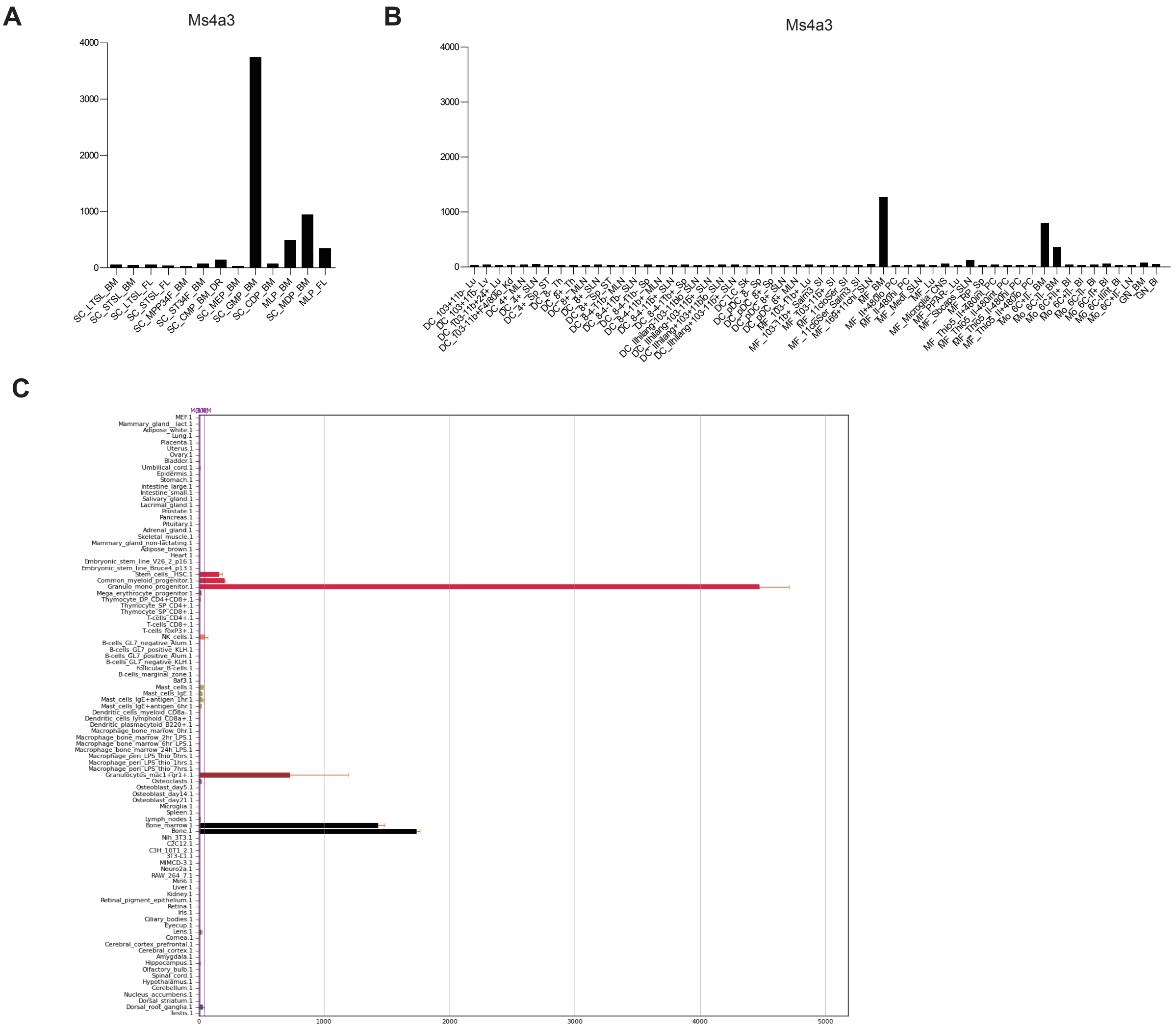

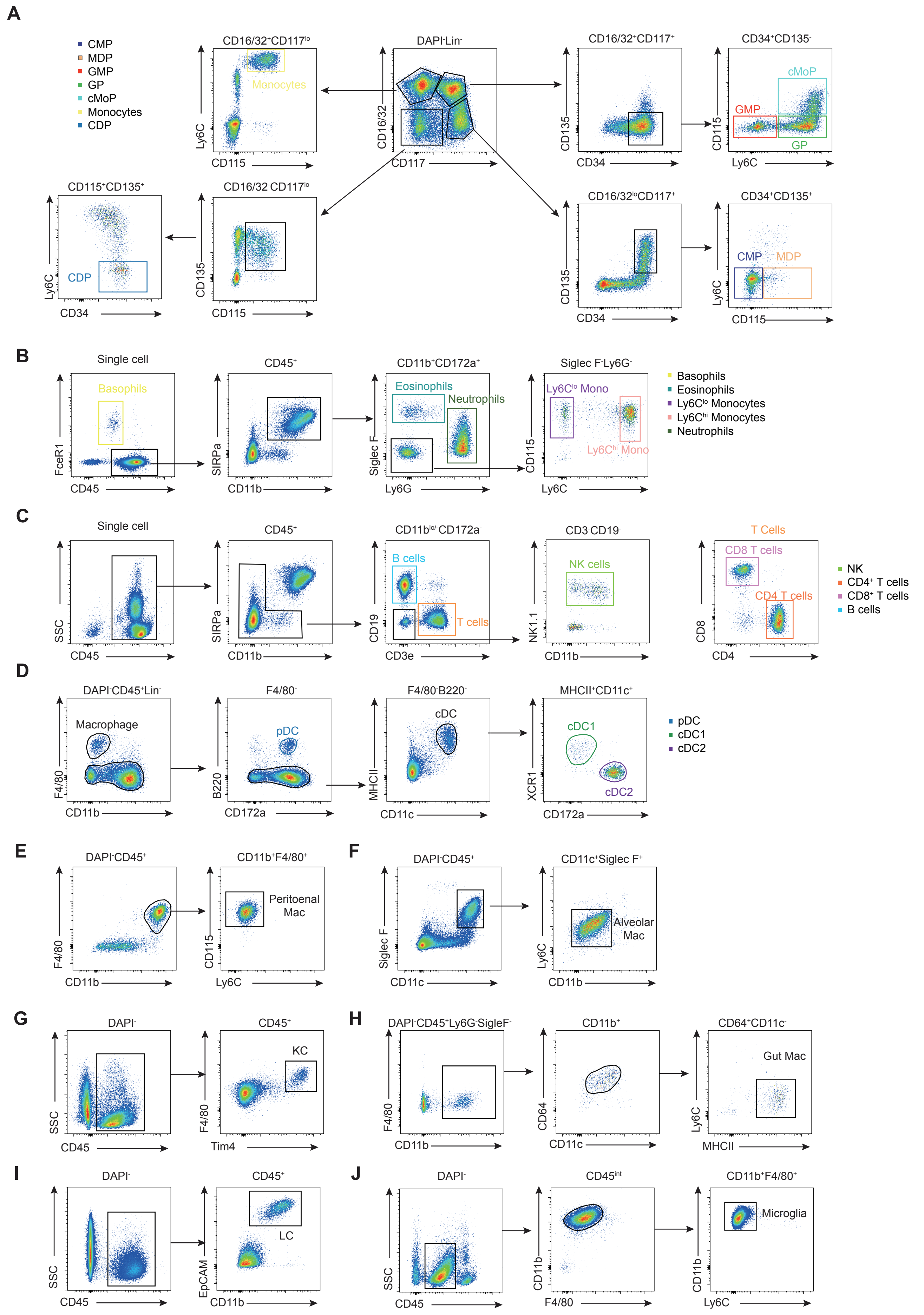

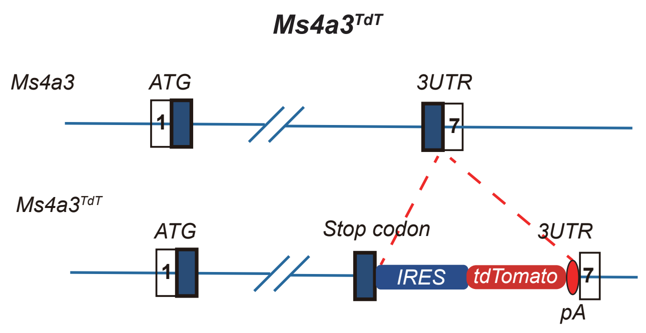

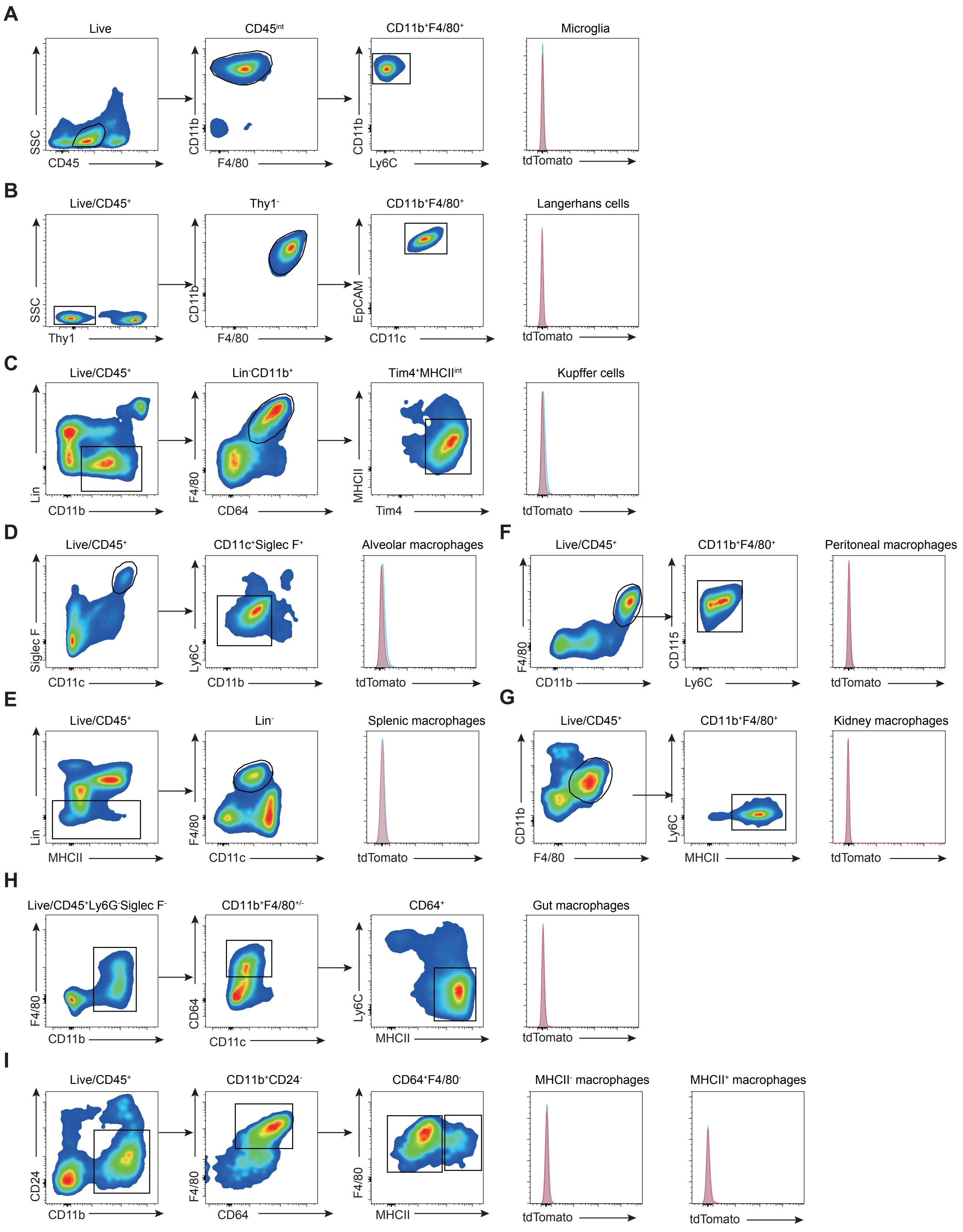

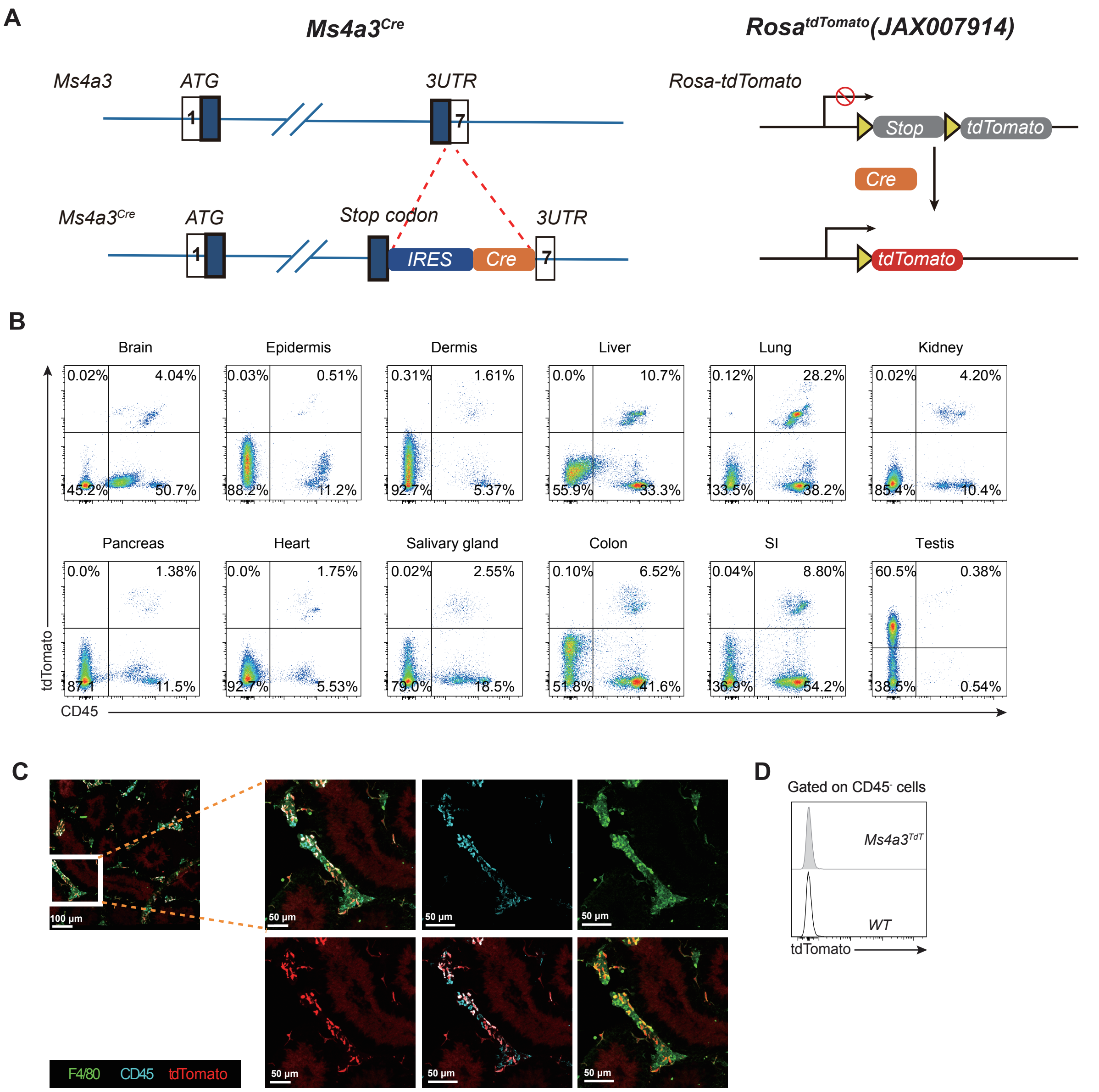

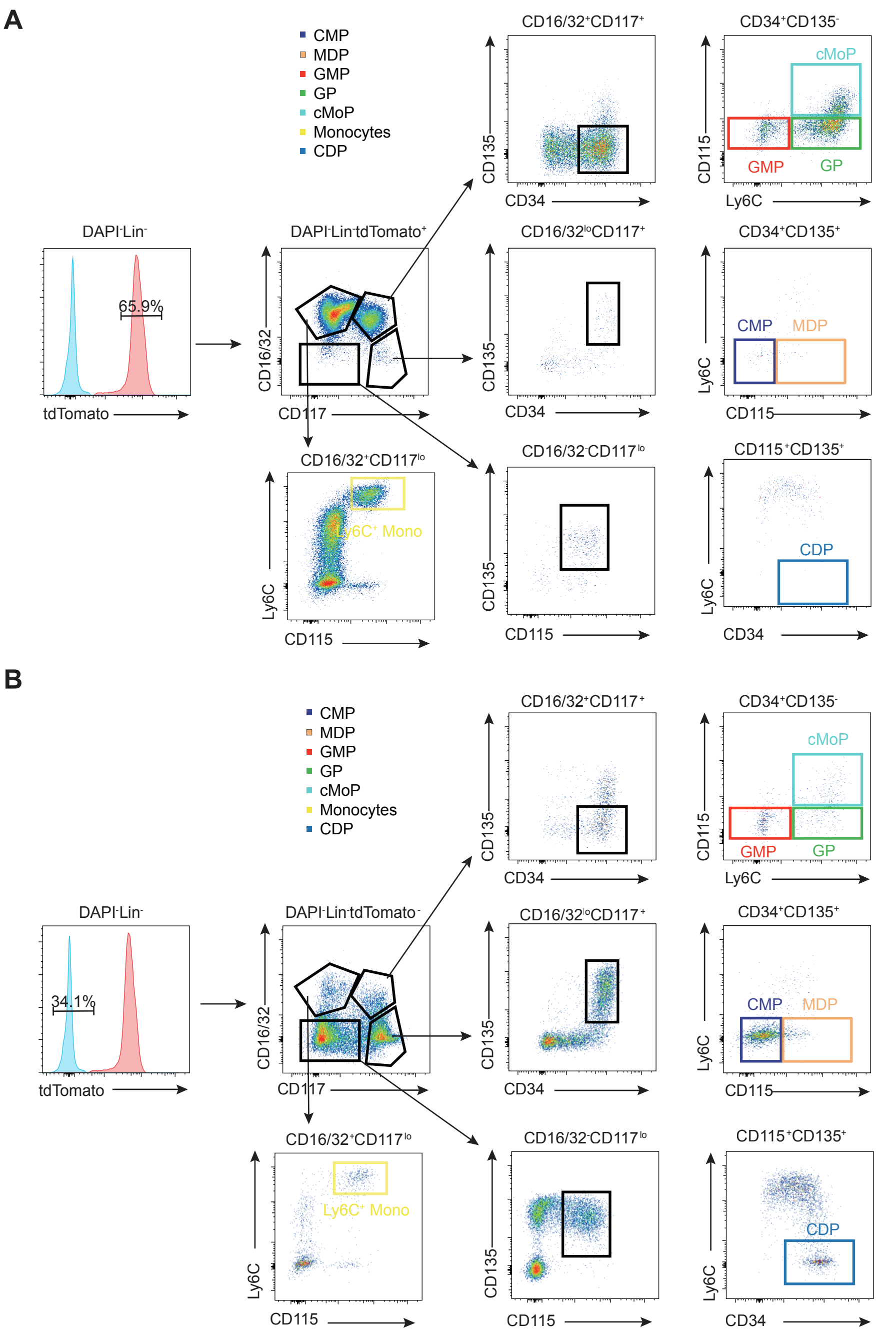

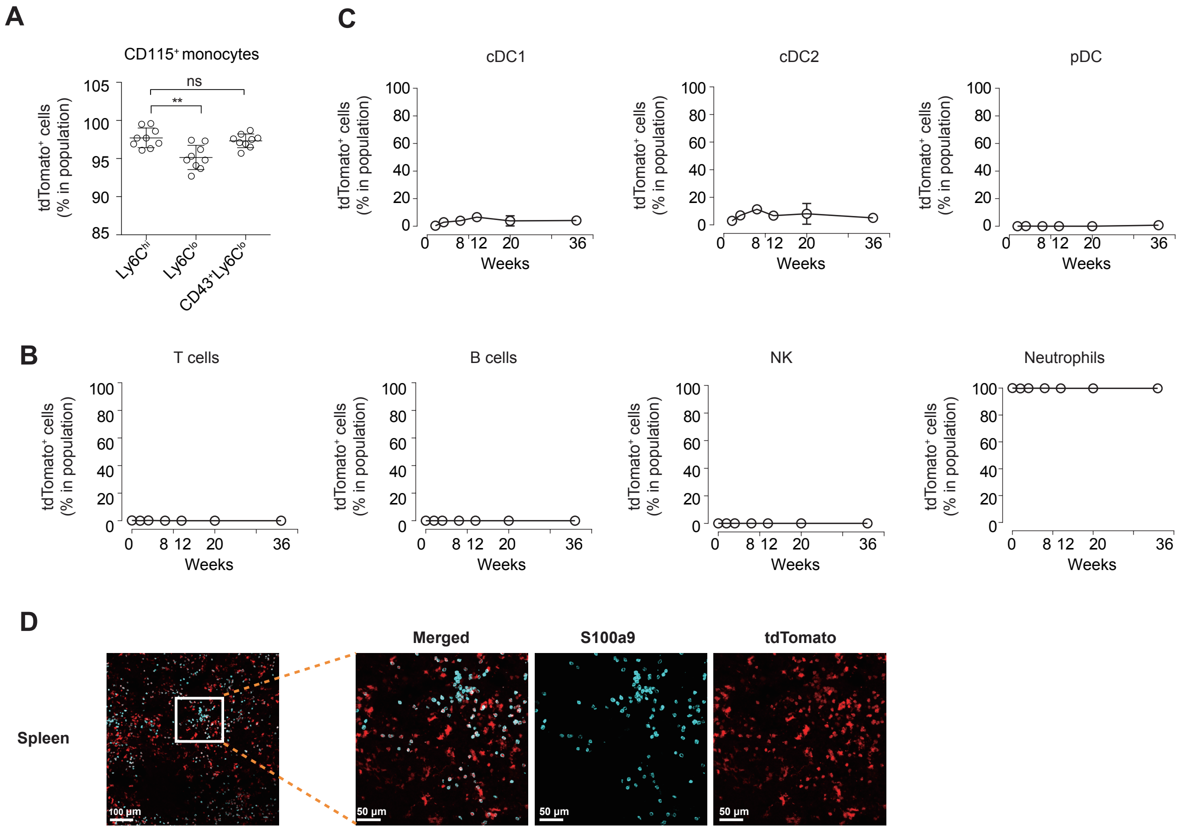

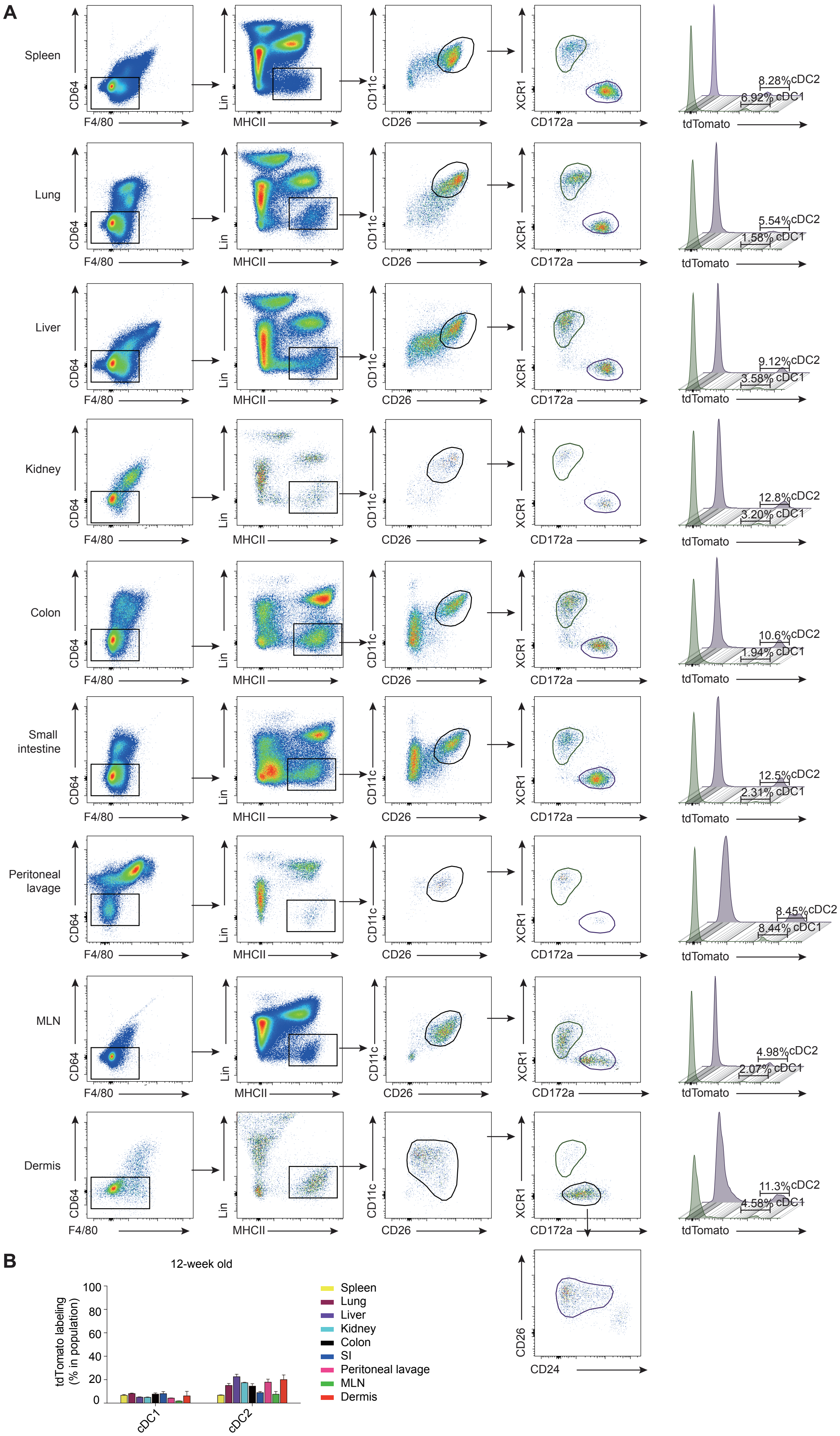

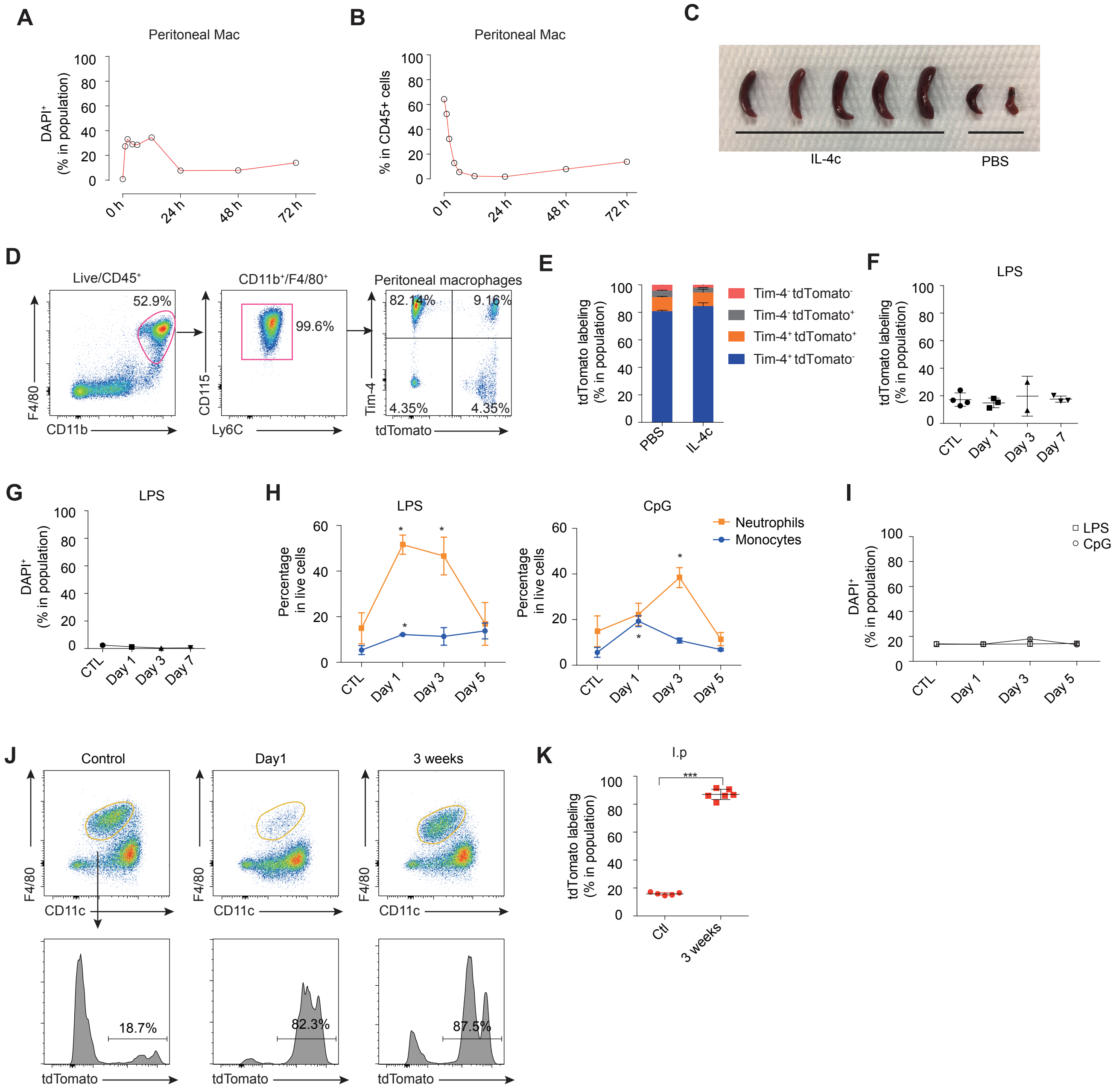
