## Supplemental Table 1 for "Fate mapping via Ms4a3 expression history traces monocyte-derived cells"

| CMap DC signature | CMap Macrophage/Monocyte signature |
|---|---|
| Adam19 | 1810011H11Rik |
| Amica1 | 4632428N05Rik |
| Anpep | 6430548M08Rik |
| Ap1s3 | A930039A15Rik |
| Ass1 | Abca1 |
| Bcl11a | Abcc3 |
| Bri3bp | Abcc5 |
| Btla | Acy1 |
| Cbfa2t3 | Apoe |
| Ccr7 | Arsg |
| Ciita | Arsk |
| Cnn2 | Asph |
| Dpp4 | Atp13a2 |
| Fgl2 | Atp6ap1 |
| Flt3 | C130050O18Rik |
| Gpr114 | C1qa |
| Gpr132 | C1qb |
| Gpr68 | C1qc |
| Gpr82 | C5ar1 |
| H2-Aa | Camk1 |
| H2-Ab1 | Cd14 |
| H2-DMb2 | Cd151 |
| H2-Eb1 | Cd164 |
| H2-Eb2 | Cd302 |
| Haao | Cd33 |
| Hmgn3 | Cebpb |
| Jak2 | Cmklr1 |
| Kit | Comt |
| Klri1 | Csf3r |
| Kmo | Ctsd |
| Napsa | Ctsf |
| P2ry10 | Ctsl |
| Pstpip1 | Dhrs3 |
| Pvrl1 | Dnase2a |
| Rab30 | Dok3 |
| Runx3 | Emr1 |
| Sept6 | Engase |
| Slamf7 | Fcgr1 |
| Spint2 | Fcgr4 |
| Tbc1d8 | Fert2 |
| Traf1 | Fez2 |
| Zbtb46 | Fgd4 |
| | Fpr1 |
| | Glul |
| | Lilra5 |
| | Gpr160 |
| | Fcgr3 |

| | |
|---|---|
| | Gbp6 |
| | Wls |
| | Gpr77 |
| | Hgf |
| | Hgsnat |
| | Hmox1 |
| | Il1a |
| | Itga9 |
| | Itgb5 |
| | Klra2 |
| | Lamp2 |
| | Lonrf3 |
| | Lpl |
| | Lrp1 |
| | Ltc4s |
| | Lyplal1 |
| | Mafb |
| | Man2b2 |
| | Mavs |
| | MerTK |
| | Mgst1 |
| | Mitf |
| | Mr1 |
| | Myo7a |
| | Nr1d1 |
| | P2ry13 |
| | Pcyox1 |
| | Pecr |
| | Pilra |
| | Pilrb1 |
| | Pilrb2 |
| | Pla2g15 |
| | Pla2g4a |
| | Pld1 |
| | Pld3 |
| | Plod1 |
| | Plod3 |
| | Plxnb2 |
| | Pon3 |
| | Pros1 |
| | Pstpip2 |
| | Ptgs1 |
| | Ptplad2 |
| | Ptprm |
| | Rhob |
| | Rhoq |
| | Rnasel |
| | Rnf13 |

| | |
|---|---|
| | Sepn1 |
| | Sepp1 |
| | Serpinb6a |
| | Sesn1 |
| | Siglece |
| | Slc11a1 |
| | Slc15a3 |
| | Slc16a10 |
| | Slc16a7 |
| | Slc17a5 |
| | Slc29a1 |
| | Slc38a6 |
| | Slc38a7 |
| | Slc7a2 |
| | Slco2b1 |
| | Slpi |
| | Snx24 |
| | Sqrdl |
| | St7 |
| | Tanc2 |
| | Tbxas1 |
| | Tcn2 |
| | Tfpi |
| | Tgfbr2 |
| | Timp2 |
| | Tlr4 |
| | Tlr7 |
| | Tlr8 |
| | Tmem195 |
| | Dram2 |
| | Tmem86a |
| | Tnfrsf21 |
| | Tom1 |
| | Tpp1 |
| | Tspan14 |
| | Tspan4 |
| | Xrcc5 |
